# Vocalisations indicate behavioural type in *Glossophagine* bats

**DOI:** 10.1101/2024.09.16.613248

**Authors:** Theresa Schabacker, Raffaella Castiglione, Lysanne Snijders, Mirjam Knoernschild

## Abstract

Vocalisations play a crucial role in the social systems of many animals, and may inadvertently reveal behavioural characteristics of the sender. Bats, the second largest mammalian order, rely extensively on vocalisations due to their nocturnal lifestyle and complex social systems, making them ideal for studying links between vocalisations and consistent behavioural traits. In this study, we developed a new testing regime to investigate if consistent individual vocalisation differences in nectarivorous bats are associated with specific behavioural types. We exposed 60 wild, male *Glossophaga soricina handleyi* bats to novel and risky stressors, and assessed their behavioural and vocal responses. Proactive, exploratory, and bold bats were more likely to produce social calls, and among the vocalising bats, more agitated bats produced higher numbers of social calls. We thus show that bat vocalisation behaviour can be indicative of a certain behavioural type, potentially allowing conspecifics to assess personalities from a distance, which in turn could impact subsequent social interactions, group dynamics, and reproductive success. Our results, in combination with previous findings in birds, suggest that advertent or inadvertent long-distance broadcasting of personality may be widespread, thus opening up new exciting questions about the links between vocalisations and sociality.

## Introduction

Across taxa, vocalisations serve fundamental roles in the lives of animals [1,2], functioning in mediating mating opportunities [3], conveying information about individual qualities [4] or emotional states (reviewed in [5,6]), facilitating collective movement and group cohesion [2,7], strengthening social bonds [8], or aiding in orientation and navigation [9]. Despite its pervasive relevance, vocalisations have been notably understudied in the context of consistent between-individual differences, i.e. animal personality traits, and existing research is heavily biased towards birds. For example, in male great tits (*Parus major*), faster explorers have higher song rates before the egg-laying period [10], generally respond more strongly vocally [11], and males with higher singing activity have higher reproductive success [12]. In black-capped chick-a-dees (*Poecile atricapillus*), more exploratory birds produce more calls in stressful situations [13]. Apart from birds, vocalisations aid in assessing emotional reactivity and arousal states in farm animals and other mammals [5,6].

Bats, the second largest order of mammals, are highly vocal animals, in part due to their nocturnality and unique life histories. Bats display a remarkable variability in roosting and feeding resources. Many bat species rely on insect swarms or fruiting and flowering trees, food sources that are often highly ephemeral and occur unpredictably but highly clumped, promoting communication and eavesdropping among conspecifics [14,15]. Additionally, bats are highly social and many species live in colonies of several thousand individuals, often forming cohesive, long-term social bonds with members of their social groups [16]. All these life history traits point to the pivotal relevance of vocalisations as a key trait in a bat’s life. Bats employ two different types of vocalisations: on the one hand, echolocation calls serve the (primary) purpose of navigation and orientation in the environment and enable bats to detect, identify, and capture prey in the dark with an extraordinary high precision rate [17,18]. On the other hand, bats actively emit social vocalisations with the sole purpose of communicating information to conspecifics [19]. In both, echolocation and social vocalisations, there is extensive inter- and intra-specific variation in acoustic features, suggesting that bats can discriminate easily between con- and heterospecifics, up to the individual level [20,21]. While much research effort has focused on elucidating this between-individual variation of acoustic features, there is almost nothing known about between-individual differences in the production rates of either vocalisation. Only two exceptions exist: in a wild population of a temperate bat species, *Pipistrellus nathusii*, individuals consistently differ in ‘acoustic exploration’, a trait describing the number of echolocation calls emitted in a novel environment, indicating that individual bats vary in the degree of environmental cue sampling [22]. The neotropical bat *Thyroptera tricolor* shows individuality in social calling behaviour, with individuals consistently differing in the number of contact calls they emit to maintain group cohesion [23].

Comprehensive studies on between-individual differences in bat behaviour, not only regarding vocalisations but personality differences in general, are scarce in the scientific literature. Only around a dozen studies have examined personality differences in bats, with a notable concentration on two temperate bat species, the little and big brown bat (*Myotis lucifugus* and *Eptesicus fuscus*) [24–28]. In all existing studies, bats exhibited repeatable behaviours in exploration [24], boldness [29], aggression [25], or social tendencies [27]. In tropical nectarivorous bats, wild *Glossophaga commissarisi* displayed consistent individual behaviour in a foraging context [30]. Recent studies have revealed that the closely related *G. soricina* is a species complex with at least four distinct species (*G. antillarum, G. valens, G. soricina* and *G. handleyi*), the latter comprising two lineages (*G. mutica* and another lineage) [31]. Wild *G. s. handleyi* and captive *G. s. antillarum* show notable behavioural variations in foraging strategies and territorial behavior [32,33]. Certain individuals consistently and effectively monopolised and defended specific plants by aggressively chasing away intruders, accompanied by high-pitched chattering vocalisations. In contrast, other individuals engaged in trap-lining foraging; thereby probing and feeding on various plants along a known route [32]. This strongly suggests the presence of significant individual differences in both classical behavioural traits and vocalisation behaviour in *G. soricina*. Additionally, a thorough vocal repertoire for captive *G. s. antillarum* bats has been described and indicates a highly developed vocal communication system [34]. Given the prevalence of *G. soricina* as one of the most common bat species in Latin America and its amenability to extended housing [35], we capitalised on these factors to investigate the relationship between vocalisations and behavioural variation in *G. s. handleyi*.

In this study, our objective was to examine if vocalisations are indicative of a certain behavioural type. Concurrently, as standardised personality assessments in bats are lacking, we aimed to validate our experimental methodology. Thus, the goal of this study was threefold: (1) to demonstrate that consistent between-individual behavioural variation in bats extends beyond spatial responses and encompasses differences in vocal production levels. Therefore, we exposed individual bats repeatedly to novel and risky stressors and assessed the repeatability of spatial behaviours and social vocalisations over time and contexts, allowing us to estimate the level of variation caused by between-individual differences, i.e. personality traits. We hypothesised that bat behaviour would be individually repeatable and conform to personality traits. Moreover, (2) we explored how individual vocalisation levels relate to classic personality traits and whether there is a relationship between vocalisation behaviour and a certain behavioural type. We hypothesised that individuals would exhibit consistent differences in the level of social call production. However, we did not have a specific *a priori* hypothesis regarding which behavioural type would be more inclined to produce social calls. Additionally, we conducted control experiments (3) to validate that our measurements accurately reflect the intended traits, rather than representing artefacts from underlying behavioural processes. Conducting control experiments to validate experimental approaches has been suggested [36] but is seldom employed in animal personality studies. Here, we expected to find significant differences in averaged behaviours between test and control situations.

## METHODS

### General fieldwork

We tested 60 wild male *G. s. handleyi* bats over the course of two years (April - June 2021 and January – May 2022) in the Santa Rosa National Park, Conservation Area Guanacaste (ACG), Costa Rica (10°50ʹ21.9”N 85°37ʹ05.2”W). Work was conducted under permit no. ACG-028-2021 and ACG-PI-006-2022 issued by the Costa Rican National System of Conservation Areas (SINAC) and the Ministry for Environment and Energy (MINAE). All fieldwork complied with the laws in Costa Rica and the guidelines of Eurobats and IUCN.

### Animal capture and husbandry

Bat were captured in groups of 7 – 11 individuals each (2021: group 1: N=7, group 2: N=9, group 3: N=9; 2022: group 4: N=6, group 5: N= 8, group 6: N=10, group 7: N=11; total: N=60). We caught bats during the nighttime with mist nets (3m height x 12m length, mesh size 19mm; Ecotone, Gdynia, Poland) in the forest, close to known feeding sites, or during the day with hand nets in day roosts in a total of 7 different sites inside the Santa Rosa National Park in Sectors Santa Rosa (10°48’46.0”N 85°38’36.8”W), Horizontes (10°42’51.2”N 85°35’46.6”W) and Poco Sol (10°52’42.7”N 85°34’22.1”W). Species identification was performed with a taxonomic field key of Costa Rican bats [37]. Following species identification, we determined sex, reproductive state, forearm length (manual caliper diaMax, ± 0.1mm, Wiha Werkzeuge GmbH, Schonach, Germany), and weight (digital scale ± 0.1g, Foraco, Lotus NL B.V., The Hague, Netherlands). Bats were marked with wing punches using a biopsy punch (DCtattoo, 1.5mm Ø) for individual identification. After capture and marking, bats were housed in two large outdoor flight cages (Eureka!: Hexagon Screen House; 3.6 × 4.2 × 2.3 m, Johnson Outdoors Inc., Racine, USA). Each flight cage was equipped with one or two custom-made day roosts mounted on the ceiling and two cylindrical bird water feeders with a protruding opening. Bats were offered a diet of daily prepared nectar substitute (NektarPlus, Nekton GmbH, Pforzheim, Germany, mixing ratio 1:5 in tap water, resulting in a 17% sugar concentration) and sugar solution (organic brown sugar, Zukra, San José, Costa Rica, mixing ratio 1:5 in tap water, resulting in a 17% sugar concentration), both accessible *ad libitum* to the bats. After data acquisition was terminated (mean captivity period: 21 +/- 6 days), we released all bats at their respective site of capture.

### Experimental setup and protocol

After capture and marking, each group of bats was given an acclimatisation period of two full nights in the housing cage. After two nights of acclimatisation, experimental testing occurred on the premises of the Santa Rosa National Park. We captured bats with a hand net from the housing cage and transferred them to the experimental site (<300 m distance) in a cloth bag. Before testing, an individual’s weight was assessed, and the bat was given 2 min to acclimatise after being caught and handled. Experimental trials were conducted inside a building in an experimental flight cage (Fig 1) (Coleman: Octagon 8 family tent; 3.96 × 3.96 × 2.08 m, The Coleman Company, Inc., Chicago, USA), in which the bats could move freely. Just before the start of a trial, bats were fed with a 17% sugar solution (to mitigate the impact of immediate hunger levels on behavioural responses) before being released into the experimental arena. Trials lasted for 30 min, after which the bats were caught and transferred back into the second housing cage.

**Fig 1:**
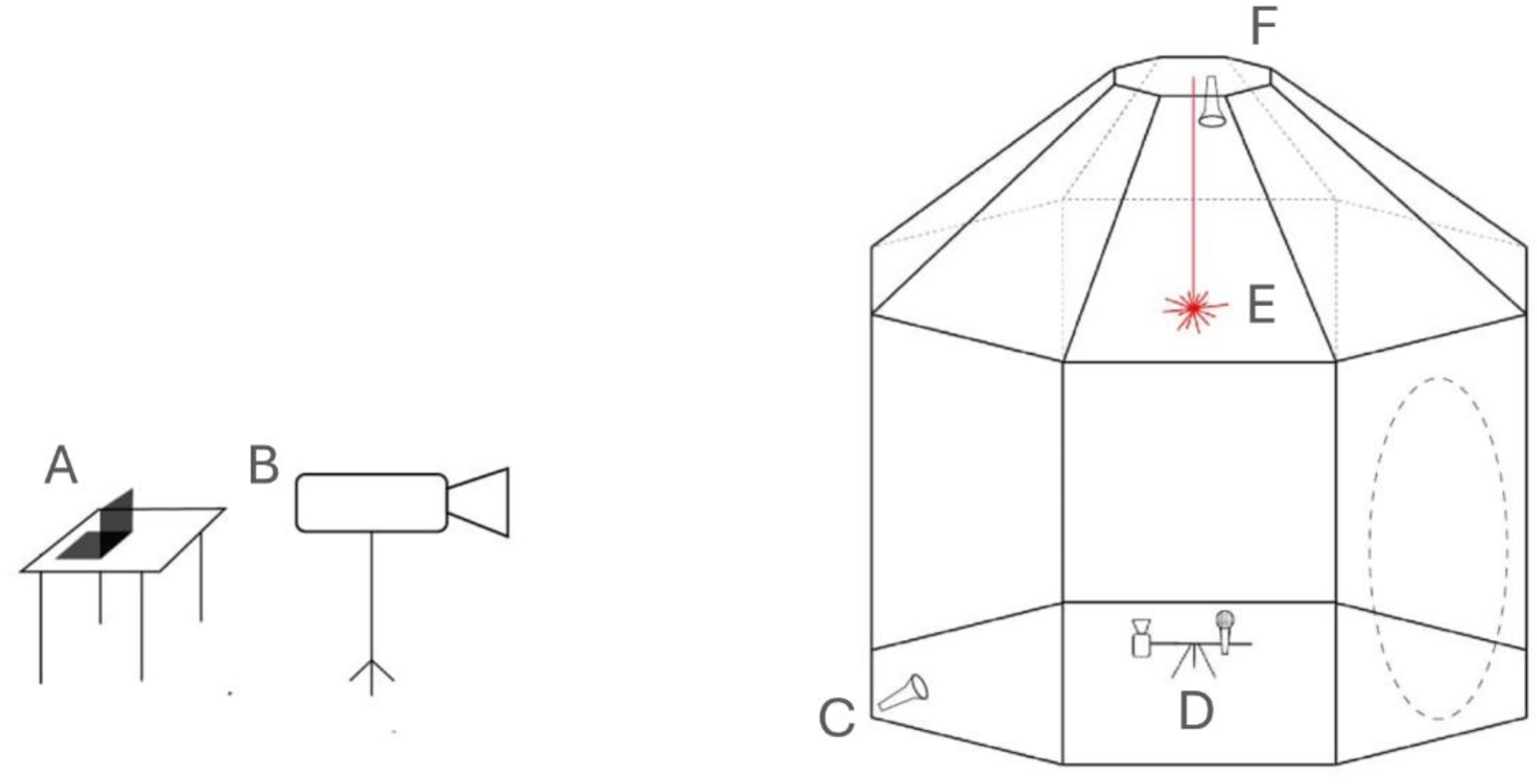
Flight cage and experimental setup for the behavioural experiments. A: Desk with portable PC running Avisoft Recorder Software. B: Overview camera. C: Infrared light source. D: Close-up camera and microphone. E: Experimental stimulus (novel/familiar object; feeder). F: Risk stimulus (white light source). © Rebecca Scheibke

All experiments were conducted at night (6.45pm – 3am) in a standardised, solitary setting. Within groups, we tested all subjects once per night in the same experiment and conducted experiments three nights in a row, followed by one night of resting. We assessed bats’ behavioural responses in three different experimental situations: Novel Environment test (NE), Novel Object test (NO), and Foraging Under Risk test (FUR), plus their respective controls: Control Novel Object test (cNO) and Control Foraging Under Risk test (cFUR). A control test for the NE test was not feasible due to the nature of the test and logistical constraints in the field. We conducted all experiments twice per group: the second trial was repeated after all bats of a respective group had performed the first trial of each experiment, resulting in an inter-trial interval of 8 +/- 2 days. The order of subjects and experiments was pseudo-randomised, with an exception for the Novel Environment test, which was always conducted first in all groups and repeated the following night (inter-trial-interval 24h, to minimise habituation effects on behavioural responses), and the control Novel Object test, which was always conducted last, to maximise habituation effects towards the familiar object (see below).

### Behavioural assays

#### Experiments

We determined individual differences in behavioural responses towards novelty-associated stress, Exploration-Avoidance, Shyness-Boldness, and the consistency thereof. The NE test is a standard open-field test and assesses an individual’s behavioural response in an unknown environment. Given that an unknown environment can offer the potential of a threat (i.e. a predator) being present, we expected this situation to induce anxiety and stress in the focal individual. Elevated spatial locomotion during a stressful situation can be interpreted as a form of active avoidance, i.e. proactive coping [38]. Therefore, we value the behaviour expressed in the NE test as indicative of an individual’s stress-coping mechanism [39]. In the remaining tests, an experimental stimulus was positioned 50 cm below the centre of the flight cage ceiling (Fig 1). In the NO test, the stimulus was a novel, echo-acoustically conspicuous object (a ball made from rubber filaments, i.e., “koosh ball”, or a hollow hemisphere, order randomised, Supplementary Fig S1). This experiment assessed the trait Exploration – Avoidance by measuring the bats’ willingness to inspect an unfamiliar object (neophilia). In the FUR test, we offered a food reward (17% nectar substitute: NektarPlus in a 1:5 ratio with tap water) alongside a risky stimulus—a 500-lumen light source from a mounted torch (2-in-1-UV-lamp, Letion, London, United Kingdom) shining a white light beam on the protruding opening of the feeder. Here, we aimed to capture variation in an individual’s willingness to move and forage in the presence of a risk stimulus (light), indicative of its inclination to engage in a risky situation [40], thus targeting the personality trait Boldness – Shyness.

#### Controls

We designed control experiments for two out of three experimental setups as suggested by Carter *et al.* [36] to ensure observed behaviours reflected the intended personality traits rather than artefacts of the experimental situation. In the cNO test, a familiar object replaced the novel object. We induced familiarity by continuously exposing the bats to this object in the housing cages, starting from the day of capture. In the cFUR test, the risk stimulus was removed, and the nectar substitute was offered without the light. An additional night of exposure to this setup was conducted before the actual cFUR test to avoid measuring responses to a novel feeding situation.

### Data capture and coding

To capture the behavioural responses of the subjects, we used two infrared-sensitive camcorders. One of the cameras (DCR-SR190, Sony, Japan) was positioned on a tripod in front of the cage, allowing for a wide overview of the experimental arena. A second camera (DCR-SR55, Sony, Japan) was positioned inside, focusing on the experimental set-up from below, allowing us to confirm approaches and interactions with the experimental stimuli. Experiments were conducted in darkness, illuminated only by infrared (IR) light (AEGIS UFLED IR lamp, Bosch, Ottobrunn, Germany). An ultrasonic microphone (Avisoft USG 116H with CM16 condenser microphone, frequency range 2–200 kHz ± 3 dB) connected to a portable PC (Lenovo S21e-20) running Avisoft-RECORDER software (R. Specht, Avisoft Bioacoustics, Glienicke, Germany; version 4.2.3102 from 16^th^ of March 2021) recorded social vocalisations.

All video recordings were coded using BORIS (Behavioral Observation Research Interactive Software [41]). For the analysis, we virtually divided the flight cage into 17 different ‘sectors’ along the walls and ceiling of the arena. In each experiment, we coded several behavioural variables related to spatial movements: e.g. the duration of the first flight after being released from the observer’s hand, the time an individual spent in flight, the number of times an individual rested or perched somewhere in the flight cage and the number of different sectors a bat visited during the experiment. Additionally, we coded behavioural variables related to interacting with the experimental stimuli: e.g. the number of times an individual inspected the unknown object, fed from or approached the feeder, and the latencies thereof (see Ethogram in Supplementary Table S1). From those coded behavioural variables, we calculated our response variables (Supplementary Table S2). All videos were coded from TS (primary observer) or RC (assistant observer). We calculated the Inter-Rater-Repeatability (IRR), to ensure consistent coding between observers. After training the assistant observer, we reached an IRR of at least 95% for each behaviour separately. All audio recordings were analysed using Avisoft SASLab Pro (R. Specht, Avisoft Bioacoustics, Glienicke, Germany). Prior to analysis, we down-sampled the sampling frequency from 500 kHz to 250 kHz and employed a high-pass filter of 10 kHz to remove noise. Spectrograms were generated using a 1024-point fast Fourier Transformation, a frame size of 100% and a Hamming window with 87.5% overlap, resulting in a bandwidth of 317 Hz and a frequency resolution of 244 Hz. Audio and video recordings were synchronised via spoken commands by the observer. We recorded individual social calls during the experiments, which resembled a specific call type classified for captive *G. s. antillarum*. Bats in captivity uttered those “alert-calls” when individuals appeared to be alert and attentive towards a situation and related to vigilant, watchful behaviours [34]. We identified and quantified the recorded social calls using Avisoft’s call detection and template-based spectrogram comparison. First, we visually inspected audio recordings and manually extracted spectrogram images of the clearest social calls to generate a template spectrogram repertoire. We then used Avisoft’s event-based option to detect sound events above a predefined amplitude threshold (threshold: 0.2V, hold-time: 0.01s). Next, we conducted a spectrogram image cross-comparisons between the detected call and reference templates from the template repertoire. We let Avisoft automatically label detected calls, which were then verified visually by the primary observer. Finally, we extracted the number of social calls produced by the focal individual for each observation.

### Statistical analysis

Due to technical difficulties and electricity failure at the field site during the conduction of the experiments, the recording of a few experimental trials failed, resulting in varying sample sizes throughout the study. A detailed sample size table can be found in Table S3 in the Supplementary Information. As a preliminary analysis, we fitted linear mixed effects models (LMM) to identify variables that significantly affected behavioural responses, using all valid test data (N_NE_=101, N_NO_=79, N_FUR_= 87). The statistical protocol for this preliminary analysis is detailed in the Method section of the Supplementary Information. Significant fixed effects for each response variable were added to the models for repeatability estimation.

Repeatability scores (R) evaluate individual variation within a population by calculating the ratio of between-individual variance (V_ind_) to the sum of V_ind_ and within-individual variance (V_e_). We used all data with two valid repeats per individual (Supplementary Table S3) and the rptR package [42] to compute adjusted repeatability scores (R_adj_) using univariate linear mixed-effects models [43]. Adjusted R scores control for confounding effects by including significant variables as fixed effects. Thus, all models were fitted with a behavioural trait as the response variable, bat identity as a random effect, and the respective significant variables assessed in the preliminary analysis as fixed effects, while always including the fixed effect ‘Trial’ to control for habituation effects. We computed confidence intervals and *p*-values for R_adj_ using parametric bootstrapping with 1,000 simulations. We evaluated model adequacy through visual examination of residual histograms, Q-Q plots, and fitted vs. residual plots. We applied a Benjamini-Hochberg procedure to correct for multiple testing. We also constructed LMM models with the lme4 package to extract variance components (V_ind_ & V_e_) for each response variable. We then averaged repeatable behavioural variables per individual across both trials, resulting in one final score per individual and experiment. This approach minimises the impact of varying external factors and extreme phenotypes [44]. Given our high repeatability scores (see Results), we believe that averaged trait scores accurately reflect the fraction of the variation attributable to individual differences, i.e., personality traits.

Next, we compared how individuals scored in the test and control situations to evaluate our methodology. We used Wilcoxon Signed Rank tests to assess differences in the mean number of object inspections between the unknown (NO) and familiar objects (cNO). Additionally, we compared the mean number of feedings between the risky (FUR) and non-risky (cFUR) situations.

To assess the relatedness of variables within tests, we conducted three separate Principal Component Analyses (PCA) on the spatial behavioural variables from each experiment. We included only trials with actively participating bats (percentage flying > 1%), resulting in sample sizes of N_NE_=57, N_NO_=45, and N_FUR_=52. We confirmed sampling adequacy with the Kaiser-Meyer-Olkin Test (KMO ≥ 0.501) and Bartlett’s Test of Sphericity (*p* < 0.001) [45]. We entered 7, 8, and 11 variables into the PCAs for the NE, NO, and FUR tests, respectively, and extracted two components each (Supplementary Table S5), as suggested by the parallel analyses. The PCA solution was Varimax-rotated and variable loadings > +/- 0.4 were considered salient (Supplementary Table S7). All subsequent analyses used individual scores from the PCA loadings.

We constructed a correlation matrix with both Principal Components (PCs) from each of the three experiments using Pearson’s correlation coefficients to assess a potential behavioural syndrome structure. A frequent caveat of behavioural syndrome research is the necessity of partitioning within- & among-individual correlations, as only the among-individual correlations constitute a behavioural syndrome, while within-individual correlations represent individual ‘correlational plasticity’ levels [46]. The state-of-the-art method is a Bayesian multivariate modelling approach, where both sources of covariation can be disentangled and analysed separately. However, Bayesian models are very data-hungry and thus inapplicable in many ecological field studies [47]. We here circumvent this caveat by using the retrieved PCs to calculate correlations. As the PCs are constructed from averaged trait values, there is no within-individual correlation present in the components. We can thus investigate the between-individual correlations (i.e. behavioural syndrome) by calculating Pearson’s correlation coefficient of the PCs. We limited our dataset to those observations where a focal individual was active during all experiments, reducing our sample size to N=43.

In the final step of the analysis, we assessed whether spatial behaviours (represented as PCs) could predict the social calling behaviour of individual bats. For each experiment separately, we fitted models with individual calling scores (averaged across trials and rounded to integers) as response variables and individual PC scores as fixed effects. Given that our dataset comprised over-dispersed count data with a high frequency of zero values (indicating no social calls recorded), we employed hurdle regression as our modelling approach, using the ‘glmmTMB’ package [48]. Hurdle models comprise two components: a binomial probability model, termed the ‘zero component,’ and a count data model, referred to as the ‘conditional component,’ which is truncated at zero values [49]. The ‘zero component’ predicts the occurrence of calling, i.e., whether the ‘hurdle’ for social calling has been surpassed. Conversely, the ‘conditional component’ predicts the count of calls when calling occurs [50], i.e. positive values reflect variability in the number of calls produced by individuals that called. For the ‘zero component,’ we employed binomial models with logit links, whereas for the ‘conditional component,’ we utilised negative binomial models with log links.

The datasets and code used in the current study are available through the Open Science Framework (OSF): https://osf.io/ewu9r/?view_only=b3799a32fadc4fdfbc8c83572edb3826

## RESULTS

Bats exhibited substantial between-individual variation in all analysed behavioural responses in the three experimental settings (see histogram and density plots for all spatial variables of each experiment in Supplementary Fig S2). Importantly, individuals differed not only in spatial but also in vocalisation behaviour. Some bats consistently produced a particular social call type (Fig 3A), formerly described as “alert call” [34]. In 104 out of 330 total trials an individual emitted at least one such social call (Fig 3B), with great variance in the number of calls produced by vocally active bats (Fig 3C).

**Fig 2:**
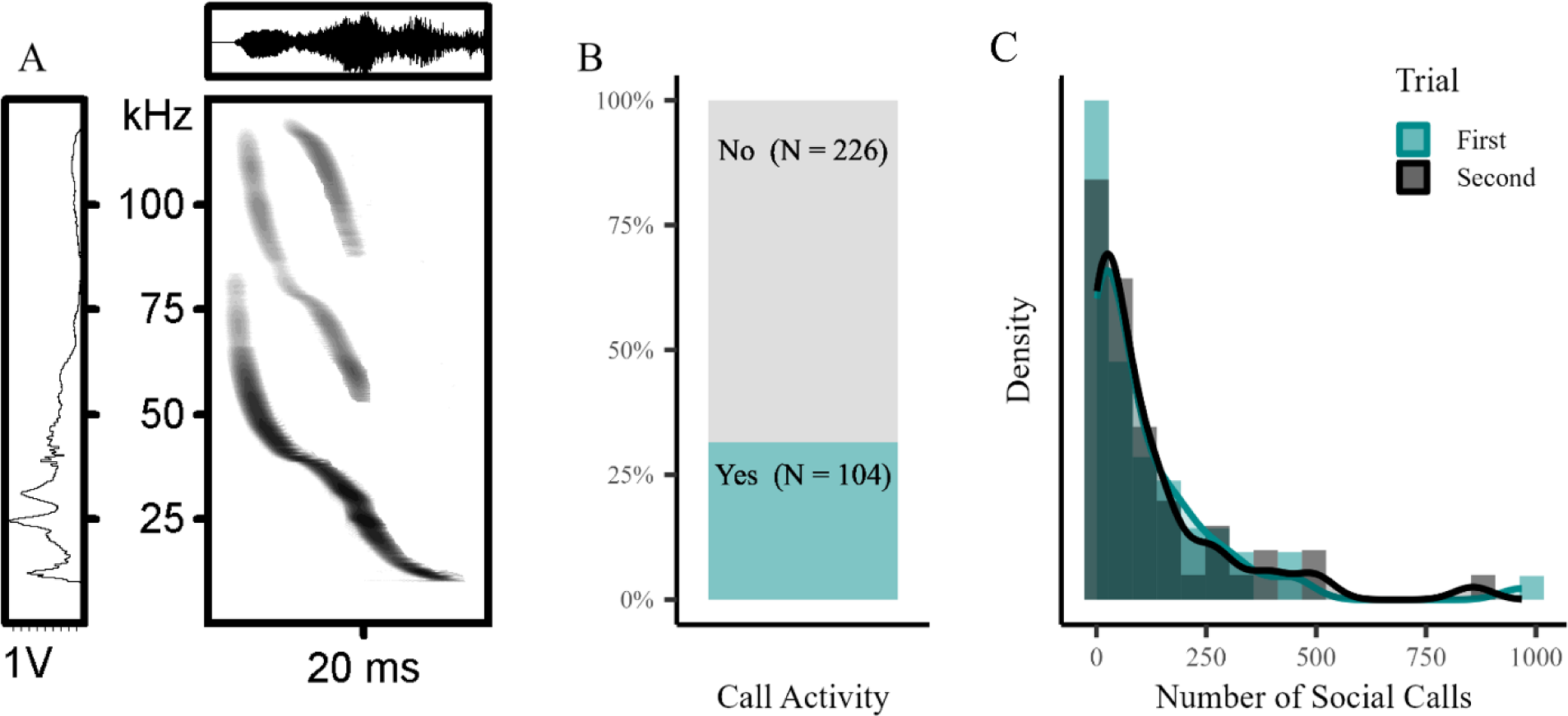
Alert calls and their prevalence among *G. s. handleyi.* (A) Oscillogram (upper panel) and spectrogram (central panel) image of an alert call, showing time on the x-axis and voltage and frequency on the y-axis, respectively. The left panel depicts the power spectrum of the call, with intensity on the x-axis and frequency on the y-axis. (B) Call activity among all bats in all trials (N=330). (C) Density and histogram plot showing the distribution of the number of social calls among the vocally active bats. Turquoise bars and line show the number of calls during the first trial, while grey bars and line depict the second trial.

**Fig 3:**
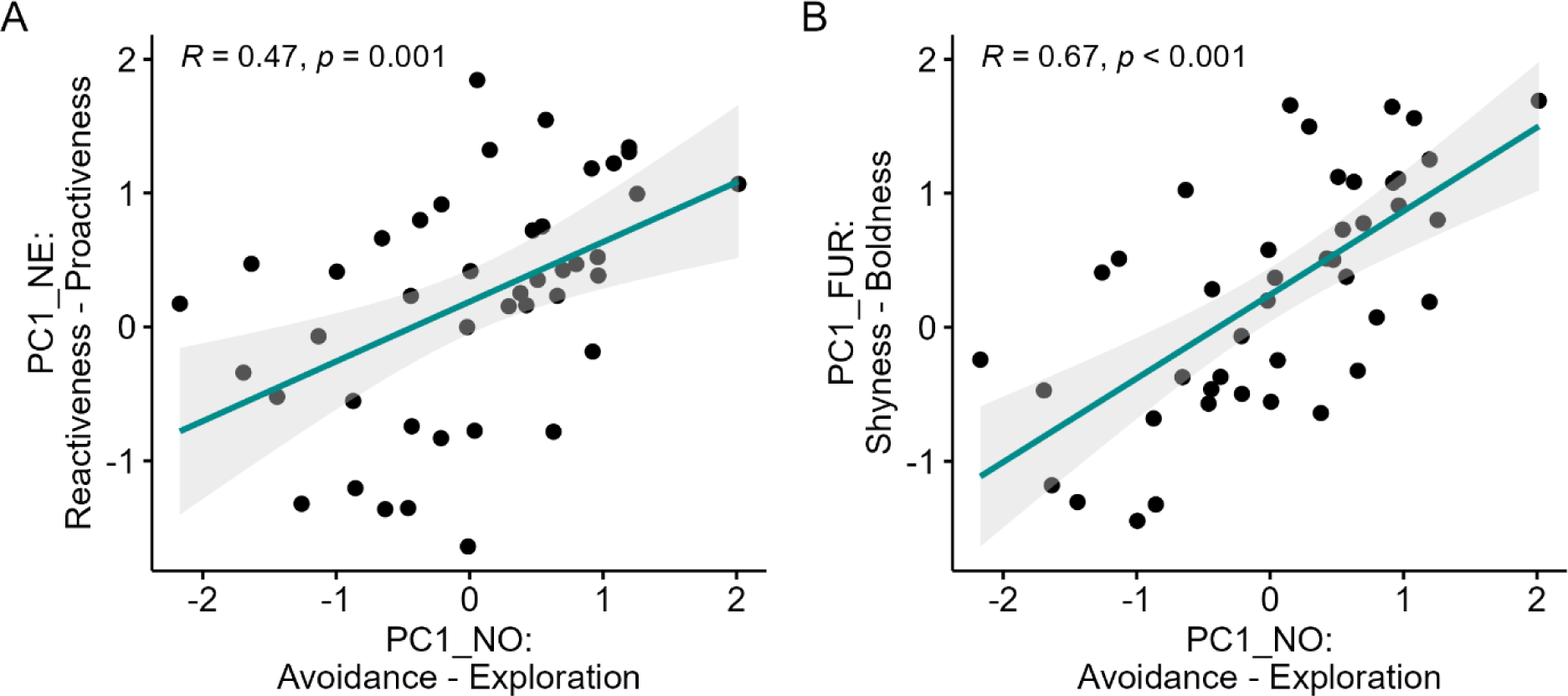
Scatterplots showing Pearson’s correlations between PC1s from different tests (N=43). Dots depict single individuals, shaded areas around the trendlines show the 95% confidence interval. (A) Strong positive correlation between Agitation - Tranquillity (NE test) and Avoidance – Exploration (NO test). (B) Strong positive correlation between Shyness – Boldness (FUR test) and Avoidance – Exploration (NO test).

### Behavioural Repeatability

Out of the 39 variables measured in the three experiments, 35 showed significant repeatability across trials, with adjusted repeatability estimates ranging from 0.31 to 0.98 (Supplementary Table S6). We also assessed cross-contextual consistency for individual spatial activity and social calling behaviour by calculating mean values across trials and calculating repeatability scores across experiments. Both spatial activity (R = 0.932, p < 0.001) and social call production (R = 0.653, p < 0.001) were highly repeatable between individuals and across experiments.

### Behavioural correlations within tests

#### Novel Environment Test

We found two principal components (PCs), which cumulatively explained 76% of the variance (Supplementary Table S7). The first axis (PC1_NE_) represented a spectrum from highly mobile individuals exploring the unknown arena to more sedentary ones and we thus interpreted it as a **Proactiveness – Reactiveness** axis. (PC1_NE_: number of perches: 0.92; percentage explored: 0.83; duration of first perch: - 0.57; duration of longest perch: -0.82; proportional variance: 43%). The second axis (PC2_NE_) represented variation in flight durations, ranging from highly restless individuals with extended flight times to more tranquil individuals with shorter flight durations. *Glossophaga soricina* is a highly active bat species [51] and expresses restlessness in elevated flight behaviour (pers. observation MK). Consequently, we interpreted this second component as an **Agitation – Tranquility** axis (PC2_NE_: duration of first flight: 0.8; duration of longest flight: 0.92; proportional variance: 33%). The variable “percentage flying” contributed significantly to both components (percentage flying PC1_NE_: 0.67; PC2_NE_: 0.68). This result becomes more intuitive when considering that the primary mode of locomotion in those bats is almost exclusively flight. Thus, both behavioural responses, i.e. how proactively/reactively individuals deal with novelty-induced stress and how agitated an individual is, were expressed as modifications of flight behaviour.

#### Novel Object Test

Two PCs cumulatively explained 68% of the variance (Supplementary Table S7). PC1_NO_ varies from actively exploring individuals, both spatially and towards the unknown object, to ‘non-explorative’ individuals, and thus we termed this axis **Exploration – Avoidance** (PC1_NO_: number of perches: 0.87; percentage explored: 0.83; percentage flying: 0.81; number of object inspections: 0.46; duration of first perch: -0.52; duration of longest perch: -0.86, proportional variance: 43%). PC2_NO_ indicated an **Agitation – Tranquillity** axis, as in the NE test ( PC2_NO_: duration of first flight: 0.82, duration of longest flight: 0.87, proportional variance: 25%).

#### Foraging Under Risk Test

Two PCs cumulatively explained 69% of the variance (Supplementary Table S7). PC1_FUR_ represented a cline from risk-prone individuals with more feedings, quicker approaches, and increased locomotion during the risky situation, to more risk-averse individuals avoiding the illuminated feeder. We thus considered it a **Boldness – Shyness** axis (number of perches: 0.85; percentage explored: 0.86; percentage flying: 0.81; number of feedings: 0.72; number of approaches: 0.8; latency to approach: -0.8; duration of first perch: - 0.57, duration of longest perch: -0.84, proportional variance: 50%). PC2_FUR_ constituted an **Agitation – Tranquillity** axis (PC2_FUR_: duration of first flight: 0.82, duration of longest flight: 0.83, proportional variance: 19%).

### Behavioural syndrome structure

We examined the relationship between PCs across different experiments to explore a potential behavioural syndrome structure in *G. s. handleyi* (Supplementary Figure S3). We found strong positive correlations between PC1_NO_ and PC1_NE_ (R = 0.47, *p* = 0.001, Fig 3A) indicating an **Exploration – Proactiveness** syndrome. Additionally, a strong positive correlation between PC1_NO_ and PC1_FUR_ (R = 0.67, *p* < 0.001, Fig 3B) indicated an **Exploration – Boldness** syndrome. Interestingly, PC1_NE_ and PC1_FUR_ were uncorrelated (R = 0.24, *p* = 0.11, Supplementary Fig S3). We found strong positive correlations between all PC2 scores (see Supplementary Figure S4), indicating consistent agitation levels between individuals across experiments.

### Does personality predict individual calling rates?

We constructed hurdle models to investigate whether personality traits (captured by PC scores) would predict individual social call production. Whether calling occurred, was significantly influenced by PC1 scores of all three experiments (Table 1, zero-inflated model). These results indicate that bats which expressed higher levels of proactiveness, exploration, and risk-taking, were more likely to emit a social call during the experimental situations. Additionally, and only in the NE test, we found a trend that more agitated bats produced more social calls in this experiment (Table 1, conditional model).

**Table 1:**
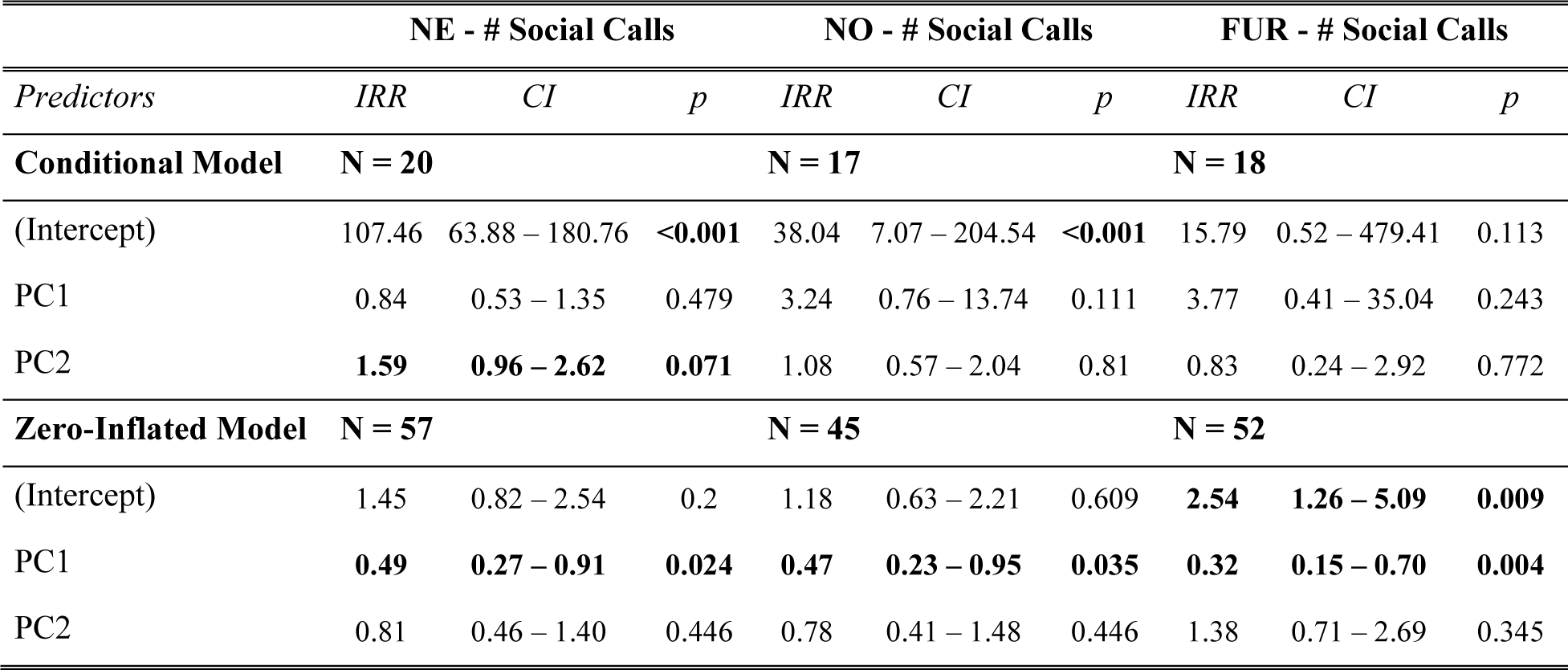
Results from hurdle models used to assess the relationship between personality traits (captured as PCs) and social calling behaviour for each experiment separately. IRR = Incidence Rate Ratios, CI = Confidence Intervalls, p = p-value. Significant values are in bold.

### Methodological Validation

As predicted, bats inspected the familiar object significantly less than the novel object (W = 693.5, *p* = 0.0025, N = 46, Fig. 4A). This suggests that the number of object inspections reflects true exploratory behaviour rather than, e.g. pure locomotion activity. Similarly, bats foraged significantly more in the non-risky setup compared to when the light was turned on (W = 111, *p* < 0.001, N = 52, Fig 4B). This indicates that the chosen stimulus (light) indeed posed a threat to the bats and the responses measured in the FUR test truly reflect risk-taking tendencies. Overall, results from control tests support our interpretation of the traits measured and validate our experimental approach.

**Fig 4:**
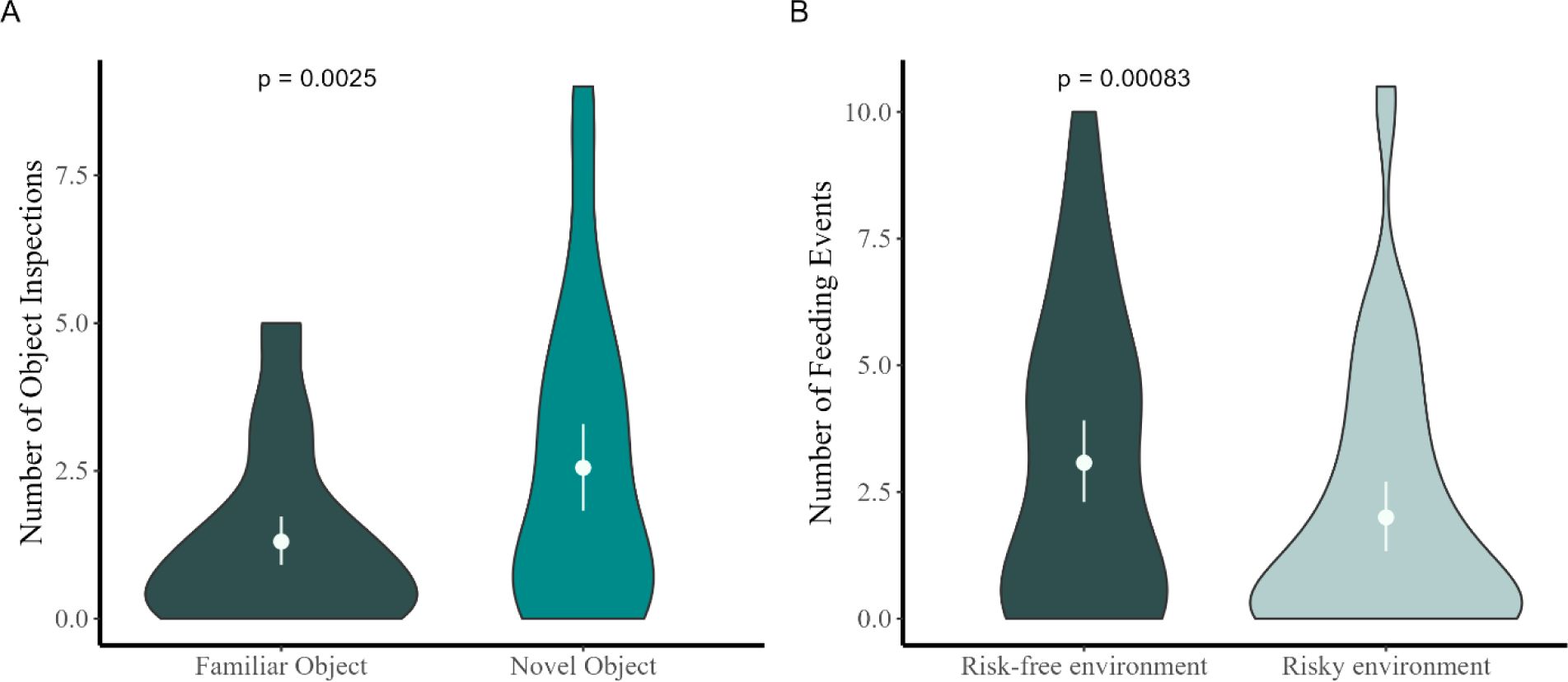
Violin plots depicting the mean number of (A) object inspections during cNO & NO (N = 46) and (B) feeding events during cFUR & FUR (N = 52). The number of object inspections and feeding events were averaged per individual across trials. The white, whiskered dots inside the violins depict the mean and 95% confidence intervals. P-values of the Wilcoxon Signed Rank Test are given in each panel for the respective test.

## DISCUSSION

Animals use vocalisations for a variety of contexts and purposes, most of which have crucial implications for sociality, reproduction, and survival [1,2], especially so in many bat species due to their nocturnality and high sociability [1,2,15]. However, that vocalisation activity may be part of an animal’s behavioural type has not gained widespread attention yet. Here, by employing a newly developed testing regime for nectarivorous bats, we reveal that *G. s. handleyi* males consistently differ in the production of a certain social call (so-called ‘alert calls’). Moreover, these calls were indicative of distinct and verified personality traits, suggesting that eavesdropping conspecifics can assess each other’s personality from a distance.

Interestingly, the more proactive, exploratory, and bold individuals demonstrated a higher likelihood of producing a social call during the experiments. These findings align with the classical definition of proactive coping: behaviours associated with a proactive coping style typically include active responses to stressors, such as aggression or exploration. These individuals tend to exhibit behaviours aimed at confronting or actively coping with challenges, rather than avoiding them [38]. Here, proactive individuals in the NE test (characterised by high levels of spatial activity) were more likely to produce social calls. The calling behaviour could thus be part of an active stress response, where individuals actively try to emit stress from their system, using both vocalisations to express their arousal and high levels of spatial locomotion, potentially searching for an escape. Conversely, highly risk-sensitive, i.e. shy, individuals might be expected to have lower thresholds for emitting alert signals in response to a stress stimulus [52]. However, we found that individuals with high boldness scores were more likely to produce alert calls, indicating that the vocalisation behaviour might indeed be an expression of a bold, i.e. risk-prone instead of risk-averse, phenotype. Some bats produced extremely high numbers of calls during the experimental situation (max number of calls: ∼1000 calls in 30 min), thereby constantly broadcasting their location. This constant dissemination of location-based information can indeed be recognised as risky behaviour *per se*. Lastly, our findings align with reports from the neotropical bat species *T. tricolor*. In this species, individuals hold a consistent role (vocal *vs.* non-vocal individuals) within their social groups [23]. Interestingly, Sagot *et al.* [53] demonstrated that an individual bat’s vocalisation behaviour predicted its exploration efforts, with more vocal bats finding more roosting opportunities in their natural habitat. Similarly, male *G. s. handelyi* bats that displayed higher levels of neophilia in the novel object test were more likely to produce an alert call. However, in our study system, the alert calls do not have a direct function in exploratory behaviour, as in *T. tricolor*, and more research is needed to fully comprehend the purpose of the social calls we recorded here.

Among the vocal bats, individuals which expressed higher levels of agitation produced more social calls during the Novel Environment test. Additionally, we noted a decline in social call production over extended periods of captivity, potentially due to a decrease in agitation caused by novel experiences like exposure to experimental conditions. The notion that acoustic signals can convey information about the agitation or arousal of the caller has been suggested in the context of within-individual variation in other bat species [54,55]. Here, considering between-individual differences, we believe a similar mechanism could be at play, where the experience of increased agitation leads to increased calling behaviour in a certain behavioural type.

If conspecifics can indeed infer another individual’s behavioural type by eavesdropping on their social vocalisations, this could influence social interactions critically. For example, Webster & Ward [56] suggest in a comprehensive review the possibility that bold individuals consistently act as leaders whereas shy individuals more often conform as followers, influencing producer – scrounger foraging tendencies within groups. If bold phenotypes broadcast their behavioural type (advertently or inadvertently) across a distance, it could lead to more scrounging behaviour among the shyer, eavesdropping conspecifics.

Consequently, the social selection pressures on specific behavioural types might shift, with the potential to affect individual fitness by altering group dynamics, foraging success and social hierarchies. However, these ideas remain speculative as we did not assess the effects on social interactions in the current experimental set-up. Thus, more research is warranted to understand the potential ecological and evolutionary implications of increased behavioural predictability of certain individuals via vocalisations. We hope that future studies take advantage of this easily observable and readily quantifiable measure and investigate how links between vocalisation and consistent behavioural differences shape social encounters, for example through vocal networks [2,57]. Additionally, it remains open to explore whether these social calls are intentionally produced signals aimed at alerting or attracting conspecifics or unintentional cues stemming from arousal or agitation.

Furthermore, it would be interesting to investigate the metabolic costs of increased social calling; Ophir *et al.* [58] demonstrated that vocalising animals may expend up to eight times more energy than silent ones. In bats, it is known that echolocation calls are extremely energetically costly, if not produced during flight [59–61]. Knowledge about metabolic costs of social calling, however, is still extremely limited. In *T. tricolor*, stationarily produced social calls increase the energetic expenditure of the vocalising bat substantially [62]. Thus, investigating potential differences in metabolic rates between individuals of *G. s. handleyi* could facilitate understanding the proximate mechanisms driving vocalisation behaviour.

## CONCLUSION

Our study revealed that consistent between-individual differences extended beyond spatial responses to vocal production rates. More exploratory, bold, and proactive bats were more likely to produce social calls in experimental settings, and among the vocal bats, more agitated individuals produced higher numbers of calls. These results suggest that vocalisation behaviour can be indicative of certain behavioural type, which has the potential to alter social interactions, group dynamics and consequently, social selection pressures.

## Acknowledgments

We thank Roger Blanco and the whole team in the Santa Rosa National Park, Costa Rica, for their support during fieldwork. We thank David Hörmann and Eva Mardus for their assistance during fieldwork. We thank Rebecca Scheibke for her illustration of the experimental setup.

