## Supplementary Information for "Vocalisations indicate behavioural type in *Glossophagine* bats"

**Research Article: “Vocalisations indicate behavioural type in Glossophagine bats”**

*Proceedings of the Royal Society B*

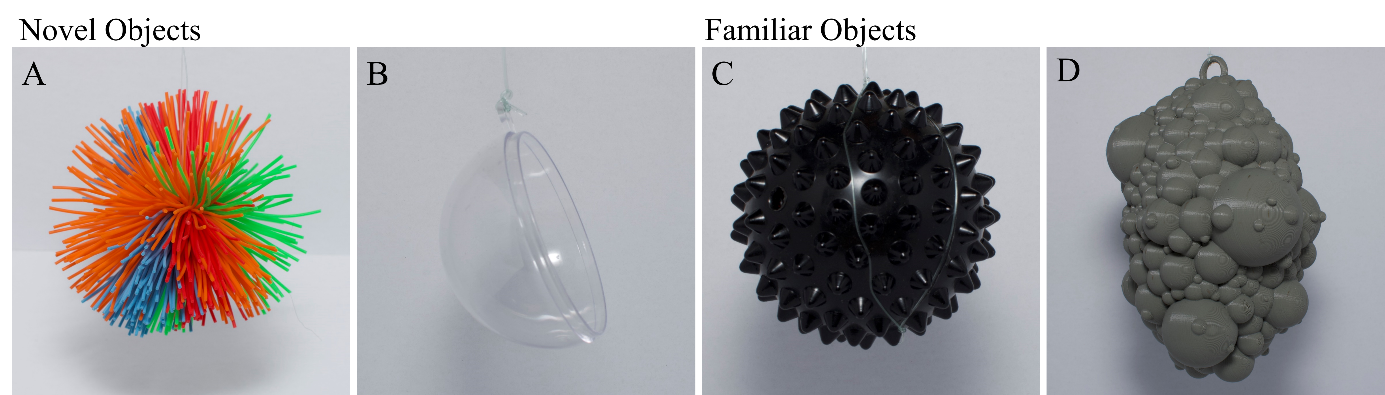
**METHODS**

Fig S1: Objects used in the NO and cNO test. A: koosh-ball, B: hollow hemisphere show objects used as *novel objects*. C: massage ball, D: 3D-printed “fake fruit” show objects used as *familiar objects*. (C) and (D) were continuously present in the housing cage starting from the day of capture.

Table S1: Ethogram used for coding of behavioural variables in Boris

| **Behavior code** | **Description** |
| --- | --- |
| Start time | Bat is released from hand |
| In flight | Bat is flying |
| Perching | Bat is hanging from one of the walls |
| Approaching | Bat is approaching/flying by, but not hovering in front of object/feeder, within one body length |
| Feeding | Bat is hovering in front of feeder opening and consuming nectar substitute |
| Hovering | Bat is levitating in front of object/feeder/light/other |
| Out of sight | Bat is not visible |
| End time | Test terminated |

| Response variable | Description | From BORIS extracted/calculated as | Assessed in test |
| --- | --- | --- | --- |
| Number of perches | Number of perches | Number of logged perches | NE, NO, FUR |
| Percentage explored | Number of unique sectors perched from | Number of sectors divided by number of sectors available (usually 12, just in the first 2 nights of testing: 9) | NE, NO, FUR |
| Percentage flying | Percentage of total time spent flying | Total flight time divided by total test duration | NE, NO, FUR |
| First flight | Duration of the first flight | Time until first perch subtracted from start time | NE, NO, FUR |
| First perch | Duration of first perch | Time from first perching until flying again | NE, NO, FUR |
| Longest flight | Duration of longest flight | Overall longest time spent flying | NE, NO, FUR |
| Longest perch | Duration of longest perch | Overall longest time spent perching | NE, NO, FUR |
| Number of object inspections | Number of times the object was inspected | Number of hoverings in front of object | NO |
| Latency to inspect | Latency till first object inspection | Time at first hovering subtracted from start time | NO |
| Number of feeding events | Number of times a bat fed from the feeder | Number of feedings | FUR |
| Latency to feed | Latency till first feeding event | Time at first feeding subtracted from start time | FUR |
| Number of approaches | Number of times a bat approached, but did not feed from the feeder | Number of approaches | FUR |
| Latency to approach | Latency till first approach | Time at first approach subtracted from start time | FUR |

Table S2: Response variables from all three experiments

Table S3: Sample Size Table

| **Sample Size Table** | | | | | |
| --- | --- | --- | --- | --- | --- |
| **N** | *NE* | *NO* | *FUR* | *Total* | *Used for:* |
| all | 112 | 109 | 109 | 330 | LMMs |
| all active | 101 | 79 | 87 | 267 | LMMs |
| repeated | 107 | 98 | 98 | 303 | rptR |
| repeated & active | 98 | 73 | 81 | 253 | rptR |
| active & averaged | 57 | 45 | 52 | 154 | PCA, Control vs Test, Hurdle Models |

Preliminary analysis – Linear Mixed Effects Models

We fitted linear mixed effects models (LMM) to identify variables that significantly affected behavioural responses, using all valid test data (N_NE_=101, N_NO_=79, N_FUR_= 87). We analysed the effect of structural factors and independent variables on the behavioural variables using the `lmer()` function from the lme4 package [1]. We included Trial (trial 1 or 2), Days in Captivity (number of days since capture; scaled covariate), Capture Method (mist net or hand net), and Weight (an individual’s weight just before the test; scaled covariate) as fixed effects, and individual ID as a random effect. We transformed non-Gaussian response variables using Tukey’s ladder of power via the `*transformTukey()*` function from the ‘rcompanion’ package [2]. We evaluated model fits by visually inspecting model residuals, residual vs. fitted plots, and Q-Q plots. For zero-inflated variables (i.e. number of object inspections, number of feedings, number of approaches, number of social calls), we ran several models with different distributional assumptions and selected the best model based on residual checks, lowest AIC, and K-fold cross-validation using the `*cv()`* function from the cv package (k = 10) [3]. We assessed collinearity using the variance inflation factor (VIF) from the ‘performance’ package [4], excluding variables with VIF > 5 and reporting those with VIF between 2.5 and 5 in Supplementary Table S4. We determined the significance of fixed effects using Type II Wald Chi-square tests, considering *p* < 0.05 as significant (Supplementary Table S4). Significant fixed effects for each response variable were added to the models for repeatability estimation.

**RESULTS**

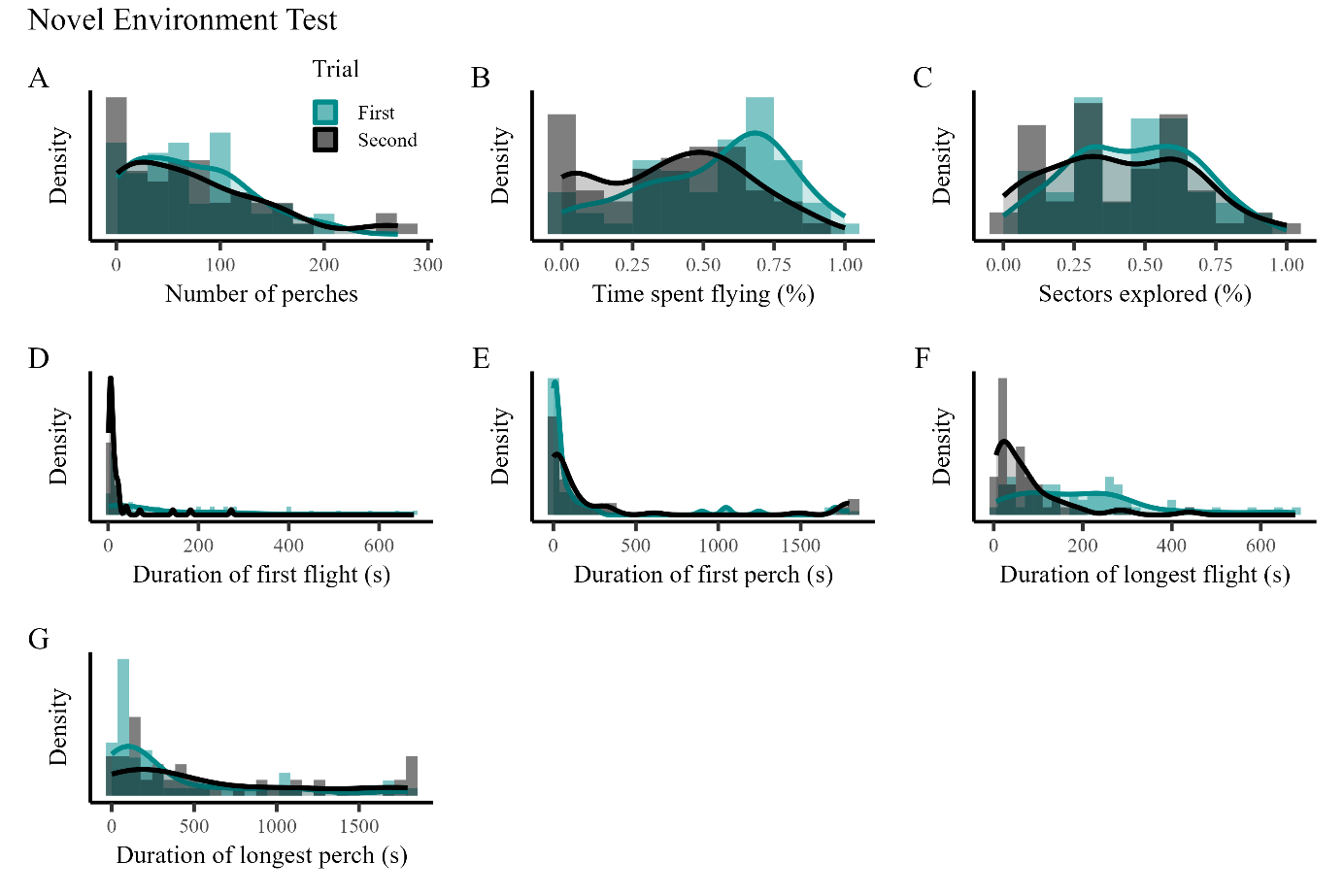
***Behavioural variation between individuals of G.s. handelyi***

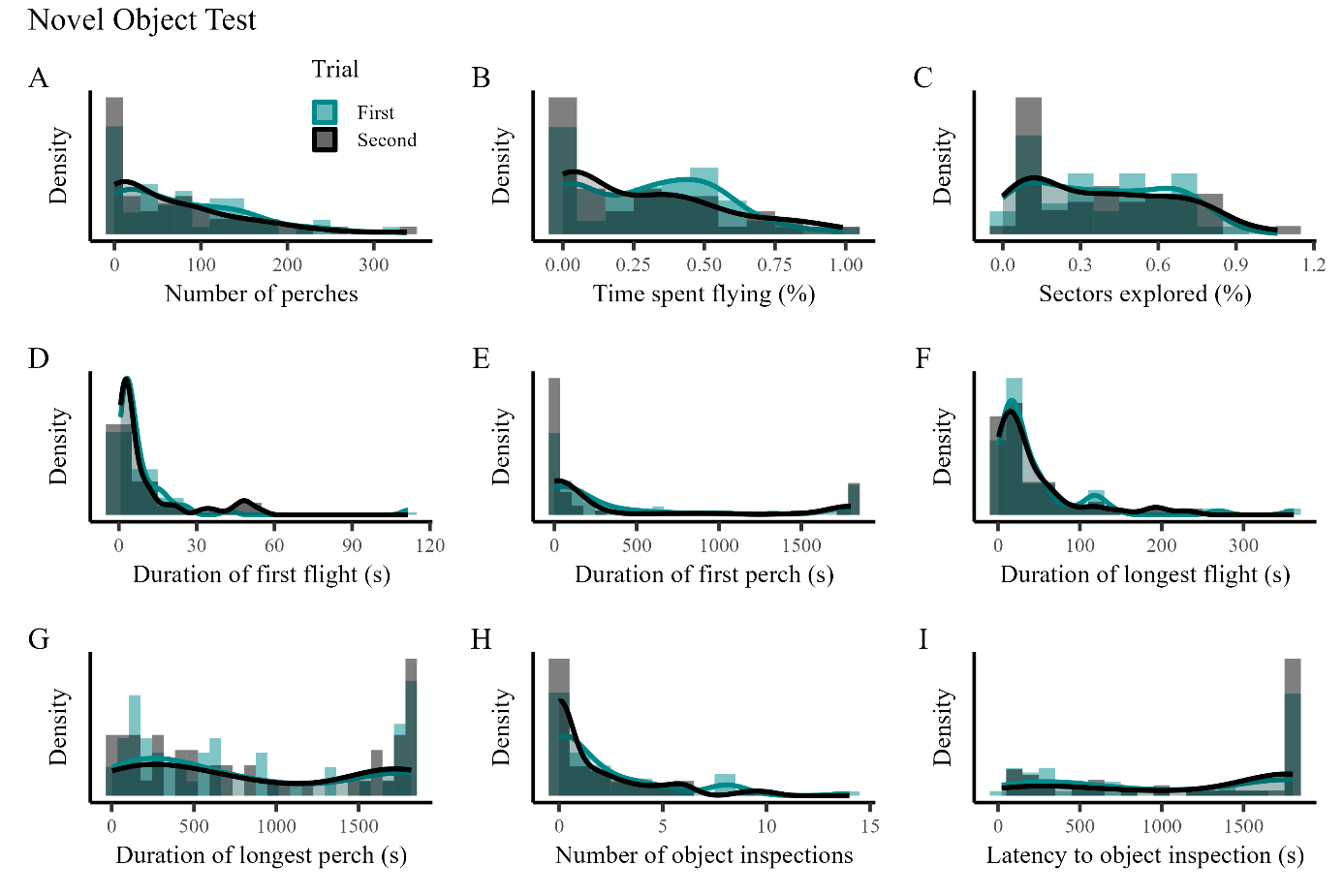

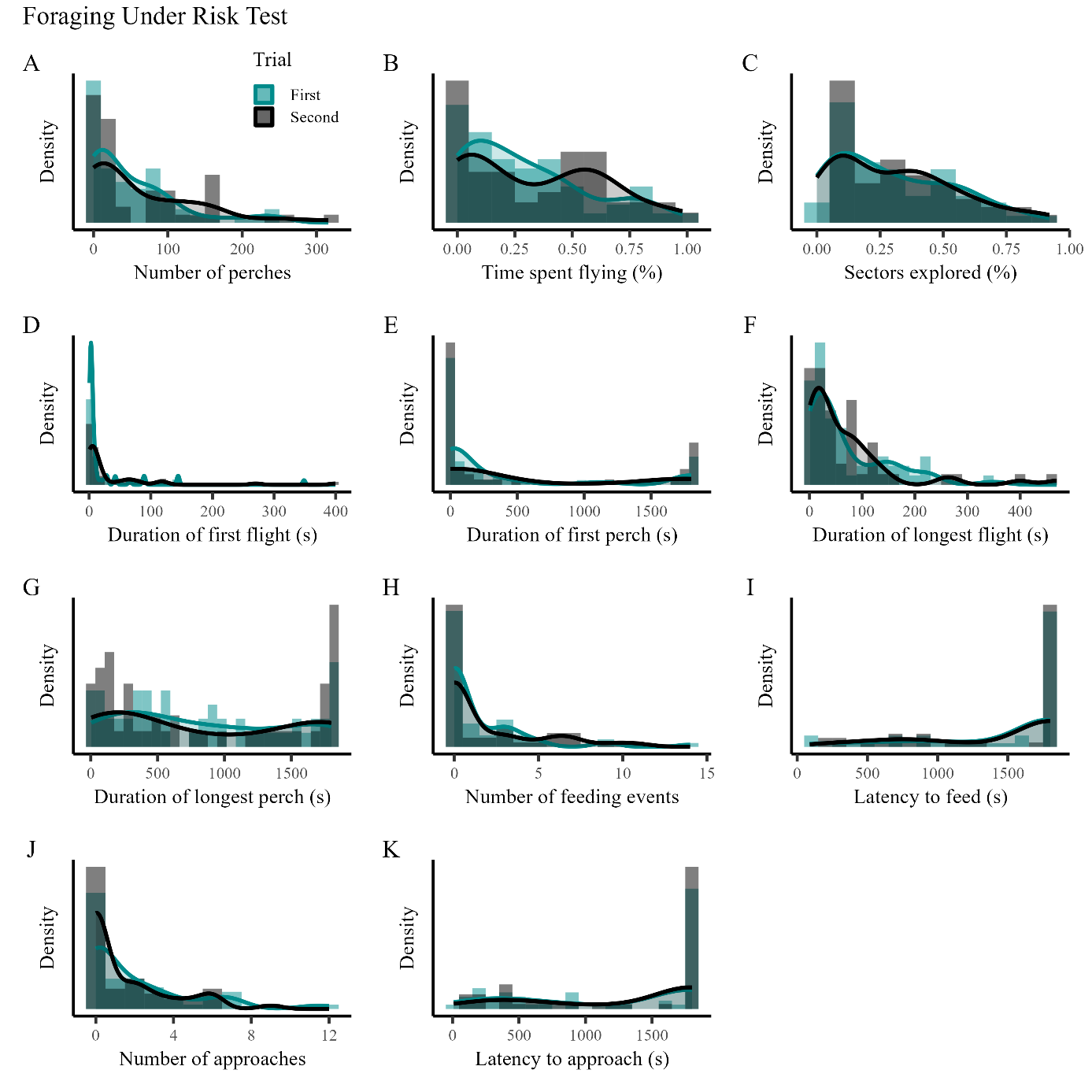
Fig S2: Histogram and density plots for all spatial variables of each experiment

| **Novel Environment Test** | | | | | | | | | |  |  |  |
| --- | --- | --- | --- | --- | --- | --- | --- | --- | --- | --- | --- | --- |
|  | **# Perches** | | | **% Flying** | | | **% Explored** | | |  |  |  |
| *Predictors* | *Est.* | *CI* | *p* | *Est.* | *CI* | *p* | *Est.* | *CI* | *p* |  |  |  |
| (Intercept) | 8.71 | 6.96 – 10.47 | **<0.001** | 0.52 | 0.42 – 0.62 | **<0.001** | 0.15 | -0.28 – 0.58 | **<0.001** |  |  |  |
| Trial [2] | 0.38 | -1.03 – 1.81 | 0.585 | **-0.11** | **-0.19 – -0.04** | **0.004** | 0.01 | -0.30 – 0.32 | 0.968 |  |  |  |
| DaysInCaptivity | 0.59 | -0.44 – 1.62 | 0.255 | 0.00 | -0.06 – 0.05 | 0.899 | -0.00 | -0.26 – 0.25 | 0.991 |  |  |  |
| CaptureMethod [Mistnet] | 0.3 | -1.84 – 2.41 | 0.789 | -0.00 | -0.14 – 0.10 | 0.764 | -0.28 | -0.81 – 0.24 | 0.284 |  |  |  |
| Weight | -0.1 | -1.08 – 0.85 | 0.817 | **-0.08** | **-0.13 – -0.03** | **0.003** | -0.14 | -0.37 – 0.08 | 0.204 |  |  |  |
| **Random Effects** | | | | | | | | | |  |  |  |
| σ^2^ | 7.60 | | | 0.02 | | | 0.29 | | |  |  |  |
| τ_00_ | 8.91 _BatID_ | | | 0.03 _BatID_ | | | 0.66 _BatID_ | | |  |  |  |
| ICC | 0.54 | | | 0.57 | | | 0.70 | | |  |  |  |
| N | 57 _BatID_ | | | 57 _BatID_ | | | 57 _BatID_ | | |  |  |  |
| Observations | 102 | | | 102 | | | 102 | | |  |  |  |
| Marginal R^2^ / Conditional R^2^ | 0.032 / 0.555 | | | 0.144 / 0.629 | | | 0.025 / 0.703 | | |  |  |  |
|  | **Duration Longest Flight** | | | **Duration Longest Perch** | | | **Duration First Flight** | | | **Duration First Perch** | | |
| *Predictors* | *Est.* | *CI* | *p* | *Est.* | *CI* | *p* | *Est.* | *CI* | *p* | *Est.* | *CI* | *p* |
| (Intercept) | 1.46 | 1.42 – 1.51 | **<0.001** | 2.41 | 2.17 – 2.66 | **<0.001** | -0.50 | -0.57 – -0.43 | **<0.001** | -0.88 | -0.91 – -0.84 | **<0.001** |
| Trial [2] | **-0.11** | **-0.14 – -0.07** | **<0.001** | 0.2 | -0.02 – 0.42 | 0.069 | **-0.16** | **-0.22 – -0.10** | **<0.001** | 0.02 | -0.02 – 0.06 | 0.429 |
| DaysInCaptivity | -0.01 | -0.03 – 0.02 | 0.57 | -0.01 | -0.15 – 0.13 | 0.901 | 0.01 | -0.03 – 0.05 | 0.649 | -0.01 | -0.03 – 0.02 | 0.574 |
| CaptureMethod [Mistnet] | -0.00 | -0.06 – 0.05 | 0.856 | 0.10 | -0.19 – 0.39 | 0.506 | 0.01 | -0.07 – 0.09 | 0.773 | 0.00 | -0.04 – 0.05 | 0.959 |
| Weight | **-0.03** | **-0.05 – -0.01** | **0.018** | **0.17** | **0.03 – 0.31** | **0.015** | -0.01 | -0.05 – 0.03 | 0.578 | 0.00 | -0.02 – 0.03 | 0.653 |
| **Random Effects** | | | | | | | | | | | | |
| σ^2^ | 0 | | | 0.20 | | | 0.01 | | | 0.01 | | |
| τ_00_ | 0.01 _BatID_ | | | 0.13 _BatID_ | | | 0.01 _BatID_ | | | 0.00 _BatID_ | | |
| ICC | 0.52 | | | 0.39 | | | 0.47 | | | 0.17 | | |
| N | 57 _BatID_ | | | 57 _BatID_ | | | 57 _BatID_ | | | 57 _BatID_ | | |
| Observations | 102 | | | 102 | | | 101 | | | 100 | | |
| Marginal R^2^ / Conditional R^2^ | 0.269 / 0.650 | | | 0.084 / 0.441 | | | 0.194 / 0.570 | | | 0.014 / 0.179 | | |

***Preliminary Analysis – Linear Mixed Effects Models***

Table S4: Results from LMMs on structural factors (fixed effects, ‘Predictors’) on behavioural variables (response variables) for all three experiments.

| **Novel Object Test** | | | | | | | | | |  |  | |  |
| --- | --- | --- | --- | --- | --- | --- | --- | --- | --- | --- | --- | --- | --- |
|  | **# Perches** | | | **% Flying** | | | **% Explored** | | |  |  | |  |
| *Predictors* | *Est.* | *CI* | *p* | *Est.* | *CI* | *p* | *Est.* | *CI* | *p* |  |  | |  |
| (Intercept) | 7.37 | 5.45 – 9.29 | **<0.001** | 0.32 | 0.20 – 0.43 | **<0.001** | 0.36 | 0.25 – 0.47 | **<0.001** |  | |  |  |
| Trial [2]* | -0.85 | -3.27 – 1.57 | 0.484 | -0.04 | -0.08 – 0.10 | 0.508 | 0.04 | -0.08 – 0.17 | 0.506 |  | |  |  |
| DaysInCaptivity* | 0.15 | -1.13 – 1.42 | 0.82 | 0.001 | -0.07 – 0.08 | 0.387 | -0.03 | -0.10 – 0.04 | 0.384 |  | |  |  |
| CaptureMethod [Mistnet] | **3.67** | **1.58 – 5.77** | **<0.001** | **0.17** | **0.04 – 0.29** | **0.013** | **0.16** | **0.03 – 0.29** | **0.013** |  | |  |  |
| Weight | **1.22** | **0.25 – 2.18** | **0.012** | **0.07** | **0.02 – 0.13** | **0.014** | 0.02 | -0.04 – 0.07 | 0.567 |  | |  |  |
| **Random Effects** | | | | | | | | | |  |  | |  |
| σ^2^ | 8.63 | | | 0.03 | | | 0.02 | | |  |  | |  |
| τ_00_ | 4.20 _BatID_ | | | 0.02 _BatID_ | | | 0.02 _BatID_ | | |  |  | |  |
| ICC | 0.33 | | | 0.39 | | | 0.56 | | |  |  | |  |
| N | 49 _BatID_ | | | 49 _BatID_ | | | 49 _BatID_ | | |  |  | |  |
| Observations | 79 | | | 79 | | | 79 | | |  |  | |  |
| Marginal R^2^ / Conditional R^2^ | 0.175 / 0.445 | | | 0.117 / 0.458 | | | 0.102 / 0.604 | | |  |  | |  |
|  | **Duration Longest Flight** | | | **Duration Longest Perch** | | | **Duration First Flight** | | | **Duration First Perch** | | | |
| *Predictors* | *Est.* | *CI* | *p* | *Est.* | *CI* | *p* | *Est.* | *CI* | *p* | *Est.* | *CI* | | *p* |
| (Intercept) | -0.58 | -0.62 – -0.54 | **<0.001** | 5.66 | 4.87 – 6.46 | **<0.001** | -0.66 | -0.76 – -0.56 | **<0.001** | 1.09 | 1.05 – 1.12 | | **<0.001** |
| Trial [2]* | -0.02 | -0.06 – 0.02 | 0.295 | 0.16 | -0.82 – 1.13 | 0.752 | 0.07 | -0.04 – 0.18 | 0.222 | -0.03 | -0.06 – 0.01 | | 0.093 |
| DaysInCaptivity* | 0.01 | -0.01 – 0.03 | 0.319 | -0.06 | -0.58 – 0.46 | 0.815 | -0.05 | -0.11 – 0.02 | 0.132 | -0.01 | -0.02 – 0.01 | | 0.585 |
| CaptureMethod [Mistnet] | 0 | -0.04 – 0.05 | 0.855 | **-1.2** | **-2.08 – -0.32** | **0.007** | 0.1 | -0.02 – 0.22 | 0.103 | 0.01 | -0.02 – 0.05 | | 0.446 |
| Weight | 0.01 | -0.01 – 0.03 | 0.578 | **-0.56** | **-0.95 – -0.16** | **0.005** | 0.03 | -0.02 – 0.08 | 0.196 | **-0.02** | **-0.03 – -0.00** | | **0.040** |
| **Random Effects** | | | | | | | | | | | | | |
| σ^2^ | 0 | | | 1.34 | | | 0.02 | | | 0 | | | |
| τ_00_ | 0.00 _BatID_ | | | 0.87 _BatID_ | | | 0.02 _BatID_ | | | 0.00 _BatID_ | | | |
| ICC | 0.66 | | | 0.39 | | | 0.59 | | | 0.7 | | | |
| N | 49 _BatID_ | | | 49 _BatID_ | | | 49 _BatID_ | | | 49 _BatID_ | | | |
| Observations | 79 | | | 79 | | | 79 | | | 79 | | | |
| Marginal R^2^ / Conditional R^2^ | 0.012 / 0.664 | | | 0.133 / 0.474 | | | 0.057 / 0.615 | | | 0.148 / 0.740 | | | |
|  | **# Object Inspections** | | | **Latency to inspect** | | |  |  |  |  |  | |  |
| *Predictors* | *Est.* | *CI* | *p* | *Est.* | *CI* | *p* |  |  |  |  |  | |  |
| (Intercept) | 0.79 | 0.22 – 1.36 | **0.008** | 4.54 | 4.04 – 5.03 | **<0.001** |  |  |  |  |  | |  |
| Trial [2]* | -0.51 | -1.14 – 0.12 | 0.105 | **1.17** | **0.53 – 1.80** | **<0.001** |  |  |  |  |  | |  |
| DaysInCaptivity* | -0.2 | -0.55 – 0.14 | 0.243 | -0.18 | -0.51 – 0.14 | 0.267 |  |  |  |  |  | |  |
| CaptureMethod [Mistnet] | **0.76** | **0.14 – 1.38** | **0.015** | **-0.75** | **-1.26 – -0.24** | **0.004** |  |  |  |  |  | |  |
| Weight | 0.26 | -0.02 – 0.54 | 0.061 | **-0.33** | **-0.57 – -0.08** | **0.007** |  |  |  |  |  | |  |
| ObjectType [2] | **0.58** | **0.06 – 1.11** | **0.027** | **-0.76** | **-1.30 – -0.22** | **0.005** |  |  |  |  |  | |  |
| **Random Effects** | | | | | | |  |  |  |  |  | |  |
| σ^2^ | 0.51 | | | 0.59 | | |  |  |  |  |  | |  |
| τ_00_ | 0.52 _BatID_ | | | 0.19 _BatID_ | | |  |  |  |  |  | |  |
| ICC | 0.5 | | | 0.24 | | |  |  |  |  |  | |  |
| N | 49 _BatID_ | | | 49 _BatID_ | | |  |  |  |  |  | |  |
| Observations | 82 | | | 82 | | |  |  |  |  |  | |  |
| Marginal R^2^ / Conditional R^2^ | 0.133 / 0.569 | | | 0.220 / 0.410 | | |  |  |  |  |  | |  |

*variables with 2.5 < VIF < 5. Interpret results with caution

| **Foraging Under Risk Test** | | | | | | | | | | | | | | | | | | | | | |
| --- | --- | --- | --- | --- | --- | --- | --- | --- | --- | --- | --- | --- | --- | --- | --- | --- | --- | --- | --- | --- | --- |
|  | | **# Perches** | | | | | | **% Flying** | | | | | | **% Explored** | | | | |  |  |  |
| *Predictors* | | *Est.* | | *CI* | | *p* | | *Est.* | | *CI* | | *p* | | *Est.* | | *CI* | | *p* |  |  |  |
| (Intercept) | | 3.05 | | 2.25 – 3.84 | | **<0.001** | | 0.55 | | 0.43 – 0.66 | | **<0.001** | | 0.41 | | 0.30 – 0.51 | | **<0.001** |  |  |  |
| Trial [2] | | 0.65 | | -0.17 – 1.46 | | 0.114 | | -0.06 | | -0.17 – 0.05 | | 0.275 | | 0.08 | | -0.04 – 0.20 | | 0.193 |  |  |  |
| DaysInCaptivity | | -0.25 | | -0.69 – 0.19 | | 0.263 | | **0.08** | | **0.02 – 0.14** | | **0.007** | | -0.04 | | -0.10 – 0.02 | | 0.212 |  |  |  |
| CaptureMethod [Mistnet] | | 0.83 | | -0.10 – 1.76 | | 0.076 | | 0.04 | | -0.10 – 0.17 | | 0.580 | | 0.09 | | -0.03 – 0.20 | | 0.153 |  |  |  |
| Weight | | 0.13 | | -0.30 – 0.57 | | 0.539 | | 0 | | -0.06 – 0.06 | | 0.968 | | 0.01 | | -0.05 – 0.07 | | 0.740 |  |  |  |
| **Random Effects** |  | |  | |  | |  | |  | |  | |  | |  | |  | |  |  |  |
| σ^2^ | | 1.07 | | | | | | 0.02 | | | | | | 0.03 | | | | |  |  |  |
| τ_00_ | | 1.47 _BatID_ | | | | | | 0.04 _BatID_ | | | | | | 0.02 _BatID_ | | | | |  |  |  |
| ICC | | 0.58 | | | | | | 0.67 | | | | | | 0.43 | | | | |  |  |  |
| N | | 55 _BatID_ | | | | | | 55 _BatID_ | | | | | | 55 _BatID_ | | | | |  |  |  |
| Observations | | 87 | | | | | | 87 | | | | | | 87 | | | | |  |  |  |
| Marginal R^2^ / Conditional R^2^ | | 0.056 / 0.604 | | | | | | 0.081 / 0.699 | | | | | | 0.040 / 0.450 | | | | |  |  |  |
|  | | **Duration Longest Flight** | | | | | | **Duration Longest Perch** | | | | | | **Duration First Flight** | | | | | **Duration First Perch** | | |
| *Predictors* | | *Est.* | | *CI* | | *p* | | *Est.* | | *CI* | | *p* | | *Est.* | | *CI* | | *p* | *Est.* | *CI* | *p* |
| (Intercept) | | -0.9 | | -0.92 – -0.89 | | **<0.001** | | 7.35 | | 6.03 – 8.67 | | **<0.001** | | -0.57 | | -0.68 – -0.46 | | **<0.001** | -0.89 | -0.95 – -0.83 | **<0.001** |
| Trial [2] | | 0 | | -0.01 – 0.01 | | 0.638 | | 0.7 | | -0.76 – 2.16 | | 0.343 | | -0.04 | | -0.15 – 0.08 | | 0.507 | 0.03 | -0.02 – 0.09 | 0.244 |
| DaysInCaptivity | | 0 | | -0.00 – 0.01 | | 0.570 | | **-0.87** | | **-1.66 – -0.09** | | **0.027** | | **0.07** | | **0.00 – 0.13** | | **0.037** | -0.03 | -0.06 – 0.00 | 0.056 |
| CaptureMethod [Mistnet] | | 0 | | -0.02 – 0.01 | | 0.623 | | -0.58 | | -2.07 – 0.92 | | 0.441 | | -0.02 | | -0.15 – 0.10 | | 0.696 | 0.01 | -0.06 – 0.08 | 0.719 |
| Weight | | 0 | | -0.01 – 0.01 | | 0.876 | | 0.04 | | -0.67 – 0.75 | | 0.909 | | 0.01 | | -0.05 – 0.07 | | 0.678 | -0.03 | -0.06 – 0.01 | 0.111 |
| **Random Effects** | | | | | | | | | | | | | | | | | | | | | |
| σ^2^ | | 0 | | | | | | 3.74 | | | | | | 0.02 | | | | | 0 | | |
| τ_00_ | | 0.00 _BatID_ | | | | | | 3.11 _BatID_ | | | | | | 0.03 _BatID_ | | | | | 0.01 _BatID_ | | |
| ICC | | 0.62 | | | | | | 0.45 | | | | | | 0.55 | | | | | 0.63 | | |
| N | | 55 _BatID_ | | | | | | 55 _BatID_ | | | | | | 55 _BatID_ | | | | | 55 _BatID_ | | |
| Observations | | 87 | | | | | | 87 | | | | | | 87 | | | | | 87 | | |
| Marginal R^2^ / Conditional R^2^ | | 0.008 / 0.628 | | | | | | 0.084 / 0.500 | | | | | | 0.055 / 0.571 | | | | | 0.071 / 0.656 | | |
|  | | **Latency to approach** | | | | | | **Latency to feed** | | | | | |  | |  | |  |  |  |  |
| *Predictors* | | *Est.* | | *CI* | | *p* | | *Est.* | | *CI* | | *p* | |  | |  | |  |  |  |  |
| (Intercept) | | 7.86 | | 6.87 – 8.84 | | **<0.001** | | 1087.22 | | 905.63 – 1268.81 | | **<0.001** | |  | |  | |  |  |  |  |
| Trial [2] | | -0.27 | | -1.38 – 0.84 | | 0.63 | | -158.73 | | -381.95 – 64.50 | | 0.161 | |  | |  | |  |  |  |  |
| DaysInCaptivity | | 0.37 | | -0.22 – 0.97 | | 0.215 | | 74.96 | | -41.85 – 191.77 | | 0.205 | |  | |  | |  |  |  |  |
| CaptureMethod [Mistnet] | | 0.09 | | -1.01 – 1.20 | | 0.866 | | -105.33 | | -299.67 – 89.01 | | 0.284 | |  | |  | |  |  |  |  |
| Weight | | 0.17 | | -0.35 – 0.70 | | 0.514 | | 60.26 | | -34.03 – 154.56 | | 0.207 | |  | |  | |  |  |  |  |
| **Random Effects** | |  | |  | |  | |  | |  | |  | |  | |  | |  |  |  |  |
| σ^2^ | | 2.22 | | | | | | 104743.28 | | | | | |  | |  | |  |  |  |  |
| τ_00_ | | 1.58 _BatID_ | | | | | | 25826.83 _BatID_ | | | | | |  | |  | |  |  |  |  |
| ICC | | 0.42 | | | | | | 0.2 | | | | | |  | |  | |  |  |  |  |
| N | | 55 _BatID_ | | | | | | 55 _BatID_ | | | | | |  | |  | |  |  |  |  |
| Observations | | 87 | | | | | | 87 | | | | | |  | |  | |  |  |  |  |
| Marginal R^2^ / Conditional R^2^ | | 0.025 / 0.430 | | | | | | 0.074 / 0.257 | | | | | |  | |  | |  |  |  |  |
|  | | **# Feedings** | | | | | | **# Approaches** | | | | | |  | |  | |  |  |  |  |
| *Predictors* | | *IRR* | | *CI* | | *p* | | *IRR* | | *CI* | | *p* | |  | |  | |  |  |  |  |
| (Intercept) | | 1.38 | | 0.50 – 3.80 | | 0.532 | | 3.02 | | 1.59 – 5.74 | | **0.001** | |  | |  | |  |  |  |  |
| Trial [2] | | 1.74 | | 0.61 – 4.92 | | 0.298 | | 0.86 | | 0.40 – 1.84 | | 0.702 | |  | |  | |  |  |  |  |
| DaysInCaptivity | | 0.81 | | 0.38 – 1.72 | | 0.579 | | 0.96 | | 0.60 – 1.56 | | 0.878 | |  | |  | |  |  |  |  |
| CaptureMethod [Mistnet] | | 1.81 | | 0.74 – 4.42 | | 0.192 | | 0.86 | | 0.44 – 1.67 | | 0.648 | |  | |  | |  |  |  |  |
| Weight | | 0.82 | | 0.51 – 1.33 | | 0.427 | | 0.89 | | 0.62 – 1.26 | | 0.498 | |  | |  | |  |  |  |  |
| **Zero-Inflated Model** | | | | | | | | | | | | | |  | |  | |  |  |  |  |
| (Intercept) | | 0.78 | | 0.42 – 1.45 | | 0.432 | | 0.47 | | 0.27 – 0.81 | | **0.006** | |  | |  | |  |  |  |  |
| **Random Effects** | | | | | | | | | | | | | |  | |  | |  |  |  |  |
| σ^2^ | | 0.42 | | | | | | 0.45 | | | | | |  | |  | |  |  |  |  |
| τ_00_ | | 0.36 _BatID_ | | | | | | 0.33 _BatID_ | | | | | |  | |  | |  |  |  |  |
| ICC | | 0.46 | | | | | | 0.42 | | | | | |  | |  | |  |  |  |  |
| N | | 55 _BatID_ | | | | | | 55 _BatID_ | | | | | |  | |  | |  |  |  |  |
| Observations | | 87 | | | | | | 87 | | | | | |  | |  | |  |  |  |  |
| Marginal R^2^ / Conditional R^2^ | | 0.190 / 0.564 | | | | | | 0.026 / 0.434 | | | | | |  | |  | |  |  |  |  |

Est. = Estimate, IRR = Incidence Rate Ratio

*variables with 2.5 < VIF < 5. Interpret results with caution

Table S5

Results from LMM on structural factors (fixed effects, ‘Predictors’) on vocal behaviour (response variables), i.e. number of social calls emitted during all three experiments.

|  | **NE - # Social Calls** | | | **NO - # Social Calls** | | | **FUR - # Social Calls** | | |
| --- | --- | --- | --- | --- | --- | --- | --- | --- | --- |
| *Predictors* | *Est.* | *CI* | *p* | *Est.* | *CI* | *p* | *Est.* | *CI* | *p* |
| (Intercept) | **1.00** | **0.32 – 1.68** | **0.004** | 0.79 | 0.14 – 1.44 | **0.018** | 0.37 | -0.02 – 0.76 | 0.065 |
| Trial [2] | **0.48** | **0.06 – 0.90** | **0.002** | 0.25 | -0.39 – 0.88 | 0.438 | 0.01 | -0.38 – 0.39 | 0.976 |
| DaysInCaptivity | **-0.40** | **-0.81 – 0.01** | **0.054** | -0.01 | -0.37 – 0.34 | 0.939 | 0.02 | -0.19 – 0.23 | 0.861 |
| CaptureMethod [Mistnet] | -0.00 | -0.84 – 0.84 | 0.998 | 0.1 | -0.69 – 0.89 | 0.807 | 0.19 | -0.28 – 0.65 | 0.428 |
| Weight | **-0.32** | **-0.61 – -0.03** | **0.027** | -0.08 | -0.40 – 0.24 | 0.634 | -0.01 | -0.23 – 0.20 | 0.893 |
| **Random Effects** | | | | | | | | | |
| σ^2^ | 0.29 | | | 0.53 | | | 0.23 | | |
| τ_00_ | 2.03 _BatID_ | | | 1.16 _BatID_ | | | 0.39 _BatID_ | | |
| ICC | 0.87 | | | 0.69 | | | 0.63 | | |
| N | 57 _BatID_ | | | 57 _BatID_ | | | 57 _BatID_ | | |
| Observations | 102 | | | 82 | | | 87 | | |
| Marginal R^2^ / Conditional R^2^ | 0.089 / 0.885 | | | 0.015 / 0.693 | | | 0.018 / 0.641 | | |

***Behavioural repeatability***

Table S6: Results from repeatability assessment of behavioural variables from all three experiments.

| *Test* | *Variable* | *R* | *95 % CI lower - upper* | *P-value* | *V_ind_ +/- SD* | *V_e_ +/- SD* | *N Observation* | *N Groups* | *LRT/Perm.* |
| --- | --- | --- | --- | --- | --- | --- | --- | --- | --- |
| **NE** | Activity (Y/N) | **0.75** | - | **0.0123** | 39.39 +/- 6.28 | - | 106 | 53 | LRT |
|  | Social Calls (Y/N) | **0.94** | - | **< 0.001** | 60.19 +/- 7.758 | - | 90 | 45 | LRT |
|  | # Social Calls | **0.87** | 0.76 - 0.92 | **0.001** | 2.48 +/- 1.58 | 0.38 +/- 0.62 | 90 | 45 | Perm. |
|  | # Perches | **0.62** | 0.39 – 0.77 | **0.001** | 10.95 +/- 3.31 | 6.67 +/- 2.58 | 90 | 45 | LRT |
|  | Percentage Explored | **0.77** | 0.61 - 0.87 | **< 0.001** | 0.037 +/- 0.19 | 0.011 +/- 0.11 | 90 | 45 | LRT |
|  | Percentage Flying | **0.62** | 0.38 - 0.78 | **0.028** | 0.036 +/- 0.19 | 0.022 +/- 0.15 | 90 | 45 | LRT |
|  | First Flight | **0.48** | 0.22 - 0.69 | **< 0.001** | 0.014 +/- 0.12 | 0.015 +/- 0.12 | 90 | 45 | LRT |
|  | First Perch | 0.24 | 0 - 0.52 | 0.059 | 0.0007 +/- 0.025 | 0.0021 +/- 0.046 | 90 | 45 | Perm. |
|  | Longest Flight | **0.53** | 0.29 - 0.71 | **0.001** | 0.0004 +/- 0.019 | 0.0003 +/- 0.019 | 90 | 45 | Perm. |
|  | Longest Perch | **0.45** | 0.18 - 0.66 | **< 0.001** | 0.17 +/- 0.42 | 0.21 +/- 0.46 | 90 | 45 | LRT |
| **NO** | Activity (Y/N) | **0.96** | - | **< 0.001** | 102 +/- 10.1 | - | 98 | 49 | LRT |
|  | Social Calls (Y/N) | **0.71** | - | **< 0.001** | 9.82 +/- 3.13 | - | 66 | 33 | LRT |
|  | # Social Calls | **0.7** | 0.47 - 0.84 | **0.001** | 1.30 +/- 1.14 | 0.55 +/- 0.74 | 66 | 33 | LRT |
|  | # of Perches | **0.43** | 0.09 - 0.68 | **0.013** | 2.32 +/- 1.52 | 3.06 +/- 1.75 | 58 | 29 | Perm. |
|  | Percentage Explored | **0.56** | 0.24 - 0.76 | **0.003** | 0.027 +/- 0.164 | 0.021 +/- 0.146 | 58 | 29 | Perm. |
|  | Percentage Flying | **0.49** | 0.16 - 0.73 | **0.008** | 0.029 +/- 0.169 | 0.029 +/- 0.171 | 58 | 29 | Perm. |
|  | Object Inspection (Y/N) | **0.51** | **-** | **0.048** | 5.23 +/- 2.29 | - | 66 | 33 | LRT |
|  | # Object Inspections | **0.37** | 0.03 - 0.63 | **0.015** | 0.36 +/- 0.60 | 0.61 +/- 0.78 | 66 | 33 | Perm. |
|  | Latency to inspect | 0.25 | 0 - 0.54 | 0.089 | 0.22 +/- 0.47 | 0.66 +/- 0.81 | 66 | 33 | Perm. |
|  | First Flight | **0.56** | 0.23 - 0.76 | **< 0.001** | 0.03 +/- 0.17 | 0.02 +/- 0.16 | 58 | 29 | LRT |
|  | First Perch | **0.65** | 0.38 - 0.83 | **< 0.001** | 0.002 +/- 0.045 | 0.001 +/- 0.032 | 58 | 29 | LRT |
|  | Longest Flight | **0.63** | 0.34 - 0.80 | **0.002** | 0.005 +/- 0.071 | 0.003 +/- 0.054 | 58 | 29 | Perm. |
|  | Longest Perch | **0.47** | 0.09 - 0.73 | **0.011** | 1.23 +/- 1.11 | 1.41 +/-1.19 | 58 | 29 | Perm. |
| **FUR** | Activity (Y/N) | 0.18 | - | 0.115 | 1.45 +/- 1.204 | - | 98 | 49 | LRT |
|  | Social Calls (Y/N) | **0.98** | - | **< 0.001** | 233.6 +/- 15.28 | - | 68 | 34 | LRT |
|  | # Social Calls | **0.68** | 0.45 - 0.83 | **< 0.001** | 0.49 +/- 0.70 | 0.23 +/- 0.48 | 68 | 34 | LRT |
|  | # of Perches | **0.57** | 0.27 - 0.76 | **0.002** | 1.50 +/- 1.22 | 1.11 +/- 1.06 | 64 | 32 | LRT |
|  | Percentage Explored | **0.34** | 0.01 - 0.65 | **< 0.001** | 0.014 +/- 0.12 | 0.025 +/- 0.16 | 64 | 32 | LRT |
|  | Percentage Flying | **0.63** | 0.36 - 0.81 | **0.001** | 0.035 +/- 0.19 | 0.02 +/- 0.14 | 64 | 32 | Perm. |
|  | Feeding (Y/N) | **0.31** | - | **0.049** | 1.82 +/- 1.35 | - | 68 | 34 | LRT |
|  | # Feedings | **0.46** | 0.17 – 0.67 | **0.002** | 0.66 +/- 0.81 | 0.77 +/- 0.88 | 68 | 34 | LRT |
|  | Latency to feed | 0.33 | 0 - 0.61 | 0.023 | 4915 +/- 70.11 | 9861 +/- 99.30 | 68 | 34 | Perm. |
|  | Approaches (Y/N) | **0.43** | *-* | **0.021** | 3.31 +/- 1.82 | - | 68 | 34 | LRT |
|  | # Approaches | **0.62** | 0.38 – 0.78 | **0.001** | 0.88 +/- 0.94 | 0.53 +/- 0.73 | 68 | 34 | Perm. |
|  | Latency to approach | **0.41** | 0.08 - 0.65 | **0.008** | 1.03 +/- 1.01 | 1.45 +/- 1.21 | 68 | 34 | Perm. |
|  | First Flight | **0.57** | 0.29 - 0.77 | **< 0.001** | 0.03 +/- 0.16 | 0.02 +/- 0.14 | 64 | 32 | LRT |
|  | First Rest | **0.68** | 0.45 - 0.83 | **0.001** | 0.02 +/- 0.14 | 0.01 +/- 0.10 | 64 | 32 | Perm. |
|  | Longest Flight | **0.62** | 0.33 - 0.78 | **0.001** | 0.001 +/- 0.022 | 0.0003 +/- 0.0174 | 64 | 32 | Perm. |
|  | Longest Perch | **0.43** | 0.10 - 0.68 | **0.005** | 1.90 +/- 1.38 | 2.51 +/- 1.58 | 64 | 32 | Perm. |

***Trait associations within tests***

Table S7: Results from PCA for each experiment separately

| *Test* | *Variable* | *PC1* | *PC2* | *Communalities* | *Complexity* |
| --- | --- | --- | --- | --- | --- |
| **NE** | # Perches | **0.92** | -0.09 | 0.86 | 1.0 |
|  | % Explored | **0.83** | -0.05 | 0.69 | 1.0 |
|  | % Flying | **0.67** | **0.68** | 0.91 | 2.0 |
|  | Duration first flight | 0.11 | **0.8** | 0.65 | 1.0 |
|  | Duration first perch | **-0.57** | -0.46 | 0.53 | 1.9 |
|  | Duration longest flight | -0.1 | **0.92** | 0.86 | 1.0 |
|  | Duration longest perch | **-0.82** | -0.38 | 0.82 | 1.4 |
|  | Proportional Variance | 0.43 | 0.33 |  |  |
| **NO** | # Perches | **0.87** | -0.23 | 0.8 | 1.1 |
|  | % Explored | **0.86** | -0.11 | 0.76 | 1.0 |
|  | % Flying | **0.81** | 0.5 | 0.9 | 1.7 |
|  | Duration first flight | 0 | **0.82** | 0.68 | 1.0 |
|  | Duration first perch | **-0.52** | -0.33 | 0.38 | 1.7 |
|  | Duration longest flight | 0.14 | **0.87** | 0.78 | 1.1 |
|  | Duration longest perch | **-0.86** | -0.33 | 0.86 | 1.3 |
|  | # Object inspections | **0.46** | 0.13 | 0.23 | 1.2 |
|  | Proportional Variance | 0.43 | 0.25 |  |  |
| **FUR** | # Perches | **0.85** | -0.19 | 0.76 | 1.1 |
|  | % Explored | **0.86** | -0.13 | 0.75 | 1.1 |
|  | % Flying | **0.8** | 0.38 | 0.78 | 1.4 |
|  | Duration first flight | -0.08 | **0.82** | 0.69 | 1.0 |
|  | Duration first perch | **-0.57** | -0.39 | 0.48 | 1.8 |
|  | Duration longest flight | 0.24 | **0.83** | 0.76 | 1.2 |
|  | Duration longest perch | **-0.84** | -0.25 | 0.77 | 1.2 |
|  | # Feedings | **0.72** | 0.18 | 0.56 | 1.1 |
|  | # Approaches | **0.8** | 0.16 | 0.67 | 1.1 |
|  | Latency to approach | **-0.8** | -0.12 | 0.65 | 1.0 |
|  | Proportional Variance | 0.5 | 0.19 |  |  |

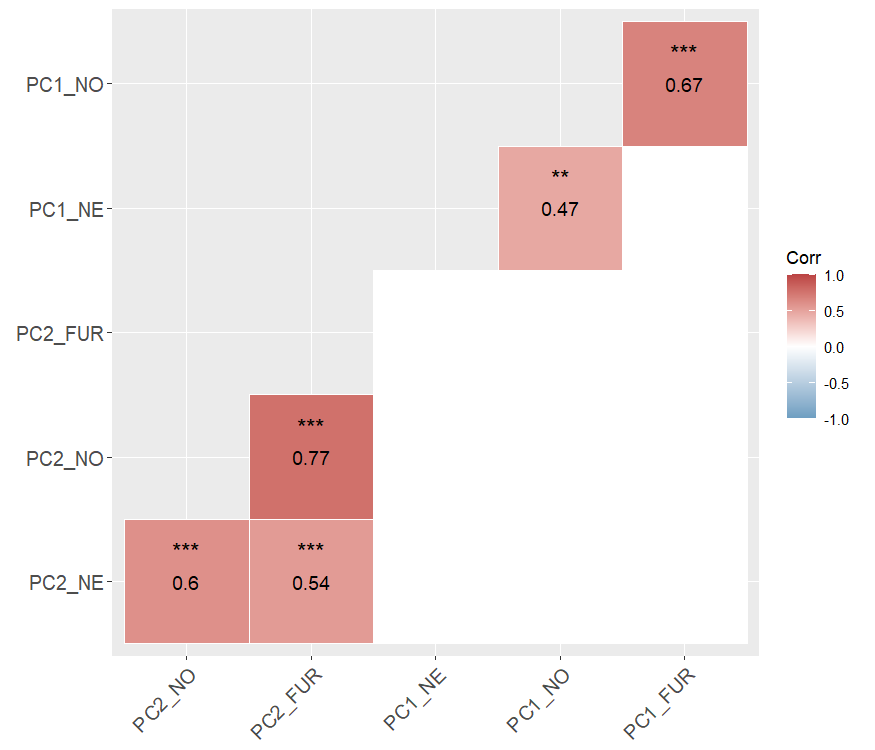
***Behavioural syndrome structure***

Fig S3: Correlation matrix showing correlations between Principle Components (PC1 and PC2) of each experiment. Red squares indicate a positive correlation, blue squares would indicate a negative relationship, blank squares indicate no significant correlation present. Numbers in the squares give Pearson’s correlation coefficient and asterisks indicate the level of statistical significance: * p < 0.05, ** p < 0.01, *** p < 0.001

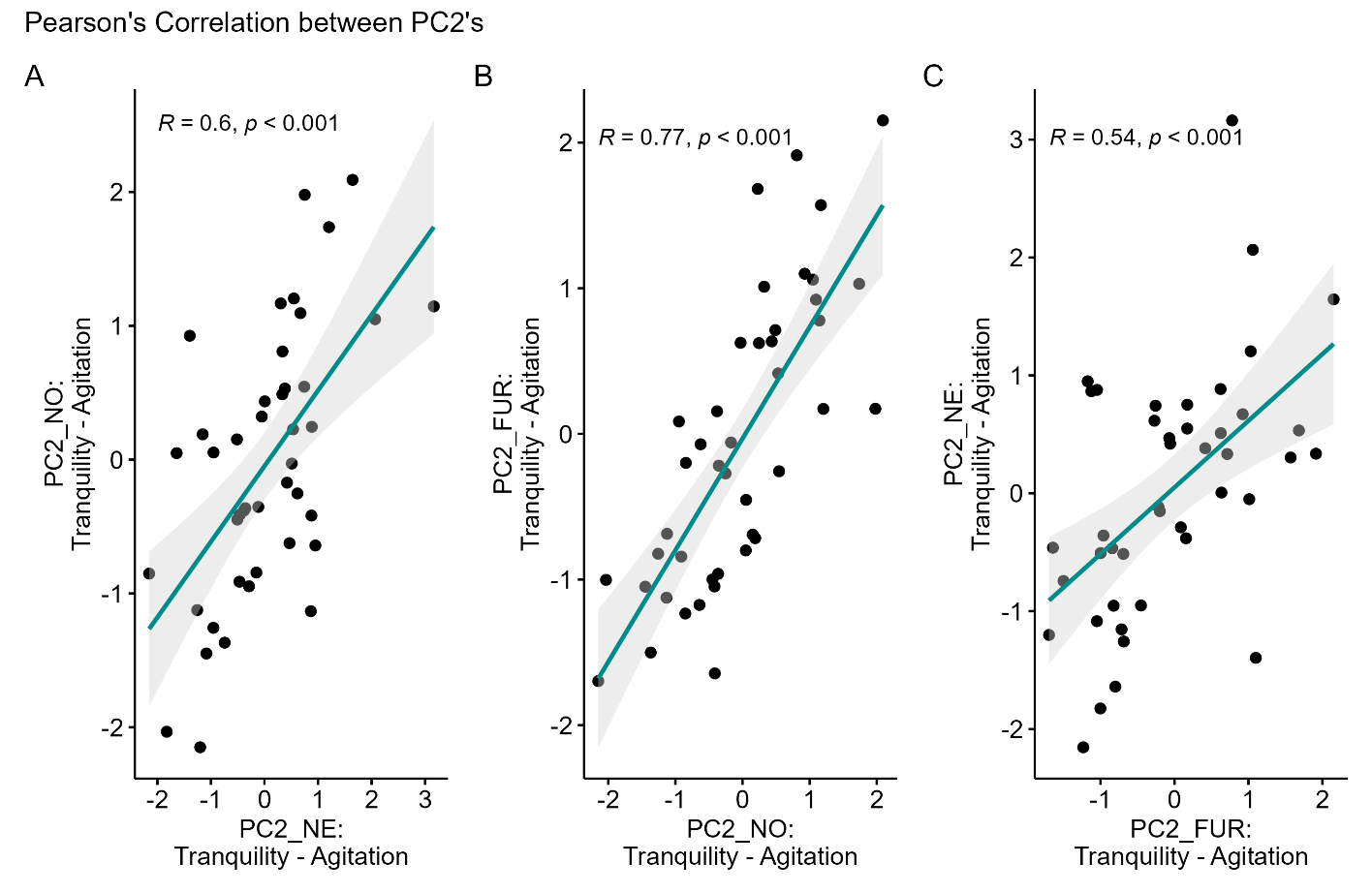
Fig S4: Scatterplots showing Pearson’s correlations between the Agitatation - Tranquility axes (PC2’s) from all three experiments (N=43). Dots depict single individuals and the shaded area around the trendline shows the 95% confidence interval. Strong positive correlations between agitation levels in (A) the NE and NO test (R=0.6, *p* < 0.001); (B) the NO and FUR test (R=0.77, *p* < 0.001); (C) the NE and FUR test (R=0.54, *p* < 0.001).
